# Linear summation of metabotropic postsynaptic potentials follows coactivation of neurogliaform interneurons

**DOI:** 10.1101/2020.12.10.418913

**Authors:** Attila Ozsvár, Gergely Komlósi, Gáspár Oláh, Judith Baka, Gábor Molnár, Gábor Tamás

**Affiliations:** MTA-SZTE Research Group for Cortical Microcircuits of the Hungarian Academy of Sciences, Department of Physiology, Anatomy and Neuroscience, University of Szeged, 52 Közép fasor, Szeged 6726, Hungary

**Keywords:** synaptic integration, neurogliaform cell, layer 1, neocortex, metabotropic GABA_B_ receptor

## Abstract

Summation of ionotropic receptor-mediated responses is critical in neuronal computation by shaping input-output characteristics of neurons. However, arithmetics of summation for metabotropic signals are not known. We characterized the combined ionotropic and metabotropic output of neocortical neurogliaform cells (NGFCs) using electrophysiological and anatomical methods. These experiments revealed that GABA receptors are activated up to 1.8 microns from release sites and confirmed coactivation of putative NGFCs in superficial cortical layers in vivo. Triple recordings from presynaptic NGFCs converging to a postsynaptic neuron revealed sublinear summation of ionotropic GABA_A_ responses and linear summation of metabotropic GABA_B_ responses. Based on a model combining distances of volume transmission from release sites and distributions of all NGFC axon terminals, we postulate that 2 to 3 NGFCs provide input to a point in the neuropil. We suggest that interactions of metabotropic GABAergic responses remain linear even if most superficial layer interneurons specialized to recruit GABA_B_ receptors are simultaneously active.

## Introduction

Each neuron in the cerebral cortex receives thousands of excitatory synaptic inputs that drive action potential output. The efficacy and timing of excitation is effectively governed by GABAergic inhibitory inputs that arrive with spatiotemporal precision onto different subcellular domains. Synchronization of GABAergic inputs appears to be crucial in structuring cellular and network excitation and behaviorally relevant rhythmic population activity (Klausberger & Somogyi, 2008). Diverse subpopulations of GABAergic neurons contribute to network mechanisms at different temporal windows and synchronized cells of particular interneuron types appear to fire in a stereotyped fashion (Klausberger & Somogyi, 2008). In general, this frequently results in coactivation of similar (and asynchronization of dissimilar) GABAergic inputs arriving to target neurons (Jang et al., 2020; Karnani et al., 2016; Kvitsiani et al., 2013), that leads to postsynaptic summation of GABAergic responses synchronously activated by presynaptic cells of the same type. Most GABAergic cell types exert inhibitory control through ionotropic GABA_A_ receptors allowing Cl^-^ ions to pass rapidly through the membrane (Barker, Ransom, & Neurobiology, 2009) and depending on the magnitude of GABA release and/or the number of synchronously active presynaptic interneurons, synaptic and extrasynaptic GABA_A_ receptors could be recruited. The integration of ionotropic inhibitory signals on the surface of target cell dendrites is temporally precise and spatially specific (Bloss et al., 2016; Klausberger, 2009; Müller, Beck, Coulter, & Remy, 2012). Summation of ionotropic receptor-mediated responses are extensively studied in the neocortex and predominantly characterized by nonlinear rules of interaction (Jadi, Polsky, Schiller, & Mel, 2012; Koch, Poggio, & Torre, 1983; London & Häusser, 2005; Qian & Sejnowski, 1990; Silver, 2010). In addition to GABA_A_ receptors, metabotropic GABA_B_ receptor activation can occur during synchronized and/or long lasting activation of GABAergic inputs (Dutar & Nicoll, 1988; Isaacson, Solis, & Nicoll, 1993; Mody, De Koninck, Otis, & Soltesz, 1994; Scanziani, 2000; Thomson & Destexhe, 1999). Among the various interneuron subtypes identified in the neocortex (Ascoli et al., 2008; Markram et al., 2004; Schuman et al., 2019), only NGFCs are known to be especially effective in recruiting metabotropic GABA_B_ receptors in addition to ionotropic GABA_A_ receptors by sporadic firing using single cell triggered volume transmission in the microcircuit (Oláh et al., 2009; Tamas, 2003). GABA binding to GABA_B_ receptors catalyzes GDP/GTP exchange at the Gα subunit and the separation of Gβγ (Bettler, Kaupmann, Mosbacher, & Gassmann, 2004). The Gβγ subunits - as membrane-anchored proteins - locally diffuse in the plasma membrane and up to four Gβγ subunits bind cooperatively to G-protein gated inward rectifier (GIRK) channels and trigger a channel opening that drives the membrane potential towards the K^+^ reverse potential (Dascal, 1997; Inanobe & Kurachi, 2014; Stanfield, Nakajima, & Nakajima, 2002; Wang, Touhara, Weir, Bean, & MacKinnon, 2016; Wickman & Clapham, 1995). Activation of GABA_B_ receptors by NGFCs control the firing of dendritic spikes in the distal dendritic domain in pyramidal cells (PCs) (Larkum, Kaiser, & Sakmann, 2002; L. M. Palmer et al., 2012; Wozny & Williams, 2011) and activity in the prefrontal cortex is effectively controlled by the strong feed-forward GABA_B_ inhibition mediated by NGFCs (Jackson, Karnani, Zemelman, Burdakov, & Lee, 2018). Moreover, GABAB receptors contribute to termination of persistent cortical activity (Craig, Mayne, Bettler, Paulsen, & McBain, 2013) and slow inhibition contributes to theta oscillations in the hippocampus (Capogna & Pearce, 2011). Relative to the summation of ionotropic responses, postsynaptic summation properties of metabotropic receptors are unexplored and to date, there has been no experimental analysis of how neurons integrate electric signals that are linked to inhibitory metabotropic receptors. We set out to test the summation of metabotropic receptor-mediated postsynaptic responses by direct measurements of convergent inputs arriving from simultaneously active NGFCs and to characterize the likelihood and arithmetics of metabotropic receptor interactions in a model of population output by incorporating experimentally determined functional and structural synaptic properties of NGFCs.

## Results

### Quantal and structural characteristics of GABAergic connections established by individual neurogliaform cells

NGFCs are capable of activating postsynaptic receptors in the vicinity of their presynaptic boutons via volume transmission (Oláh et al., 2009). To gain insight into the possible effective radius of volume transmission we characterized properties of NGFC-PC connections. *In vitro* simultaneous dual whole-cell patch clamp recordings were carried out on synaptically connected L1 NGFC to L2/3 PC pairs in brain slices from the somatosensory cortex of juvenile male Wistar rats. Pre- and postsynaptic cells were chosen based on their characteristic passive membrane and firing properties (Fig. 1A) and recorded neurons were filled with biocytin, allowing *post hoc* anatomical reconstruction of recorded cells and estimation of putative synaptic release sites (Fig. 1B, H). Single action potentials triggered in NGFCs elicit biphasic GABA_A_ and GABA_B_ receptor-mediated responses on the target neurons (Tamas, 2003). To determine the number of functional release sites, we recorded IPSCs under different release probability by varying extracellular Ca^2+^ and Mg^2+^ concentrations (Fig. 1C, F). NGFC evoked inhibitory postsynaptic potentials show robust use-dependent synaptic depression, therefore we limited the intervals of action potential triggered in NGFCs to 1 minute (Karayannis et al., 2010; Tamas, 2003). We collected a dataset of n= 8 NGFC to PC pairs with an average of 65.5 ± 5.264 trials per pair and 32.75 ± 4.155 trials for a given Mg^2+^/Ca^2+^ concentration per conditions. The limited number of trials due to the use-dependent synaptic depression of NGFCs restricted our approach to Bayesian Quantal Analysis (BQA) previously shown to be robust for the estimation of quantal parameters (Bhumbra & Beato, 2013a). As expected, IPSC peak amplitudes were modulated by elevated (3 mM) and reduced (1.5 mM or 2 mM) extracellular Ca^2+^ concentrations consistent with the decline in release probability (Fig. 1D). Distributions of IPSC amplitudes detected in paired recordings were in good agreement with the estimated quantal amplitude distribution derived from the BQA (Fig. 1E). According to BQA, the estimated mean number of functional release sites (Nfrs) was 10.96 ± 8.1 with a mean quantal size (q) of 3.93 ± 1.21 pA (Fig. 1G). Full reconstruction of functionally connected NGFC-PC pairs (n= 6) allowed comparisons of the Nfrs estimated by BQA and the number of putative release sites by counting the number of presynaptic boutons located within increasing radial distances measured from postsynaptic dendrites (Fig. 1H). Previous experiments showed that direct synaptic junctions are not required for functional NGFC to PC connections (Oláh et al., 2009) and GABA reaches receptors up to 3 μm from the release site (Farrant & Nusser, 2005; Overstreet-Wadiche & McBain, 2015; Overstreet, Jones, & Westbrook, 2000). In agreement with earlier observations (Oláh et al., 2009), direct appositions were not observed in most NGFC to PC pairs and the number of NGFC axonal boutons potentially involved in eliciting postsynaptic responses increased by systematically increasing the radial distance from the dendrites of PCs. Projecting the range of BQA-derived Nfrs estimates over the number of NGFC boutons putatively involved in transmission for the same connections (Fig. 1H, red lines) suggests an effective range of 0.86 to 1.75 μm for nonsynaptic volume transmission from NGFCs to PCs supporting previous reports on distances covered by extrasynaptic GABAergic communication (Farrant & Nusser, 2005; Overstreet-Wadiche & McBain, 2015; Overstreet et al., 2000). Moreover, we detected linear correlation (r= 0.863, p= 0.027) between BQA-derived Nfrs and the number of NGFC boutons putatively involved in transmission at radial distances <1.5 μm from PC dendrites; decreasing or increasing the distance resulted in the loss of correlation (Fig. 1I).

**Figure 1.**
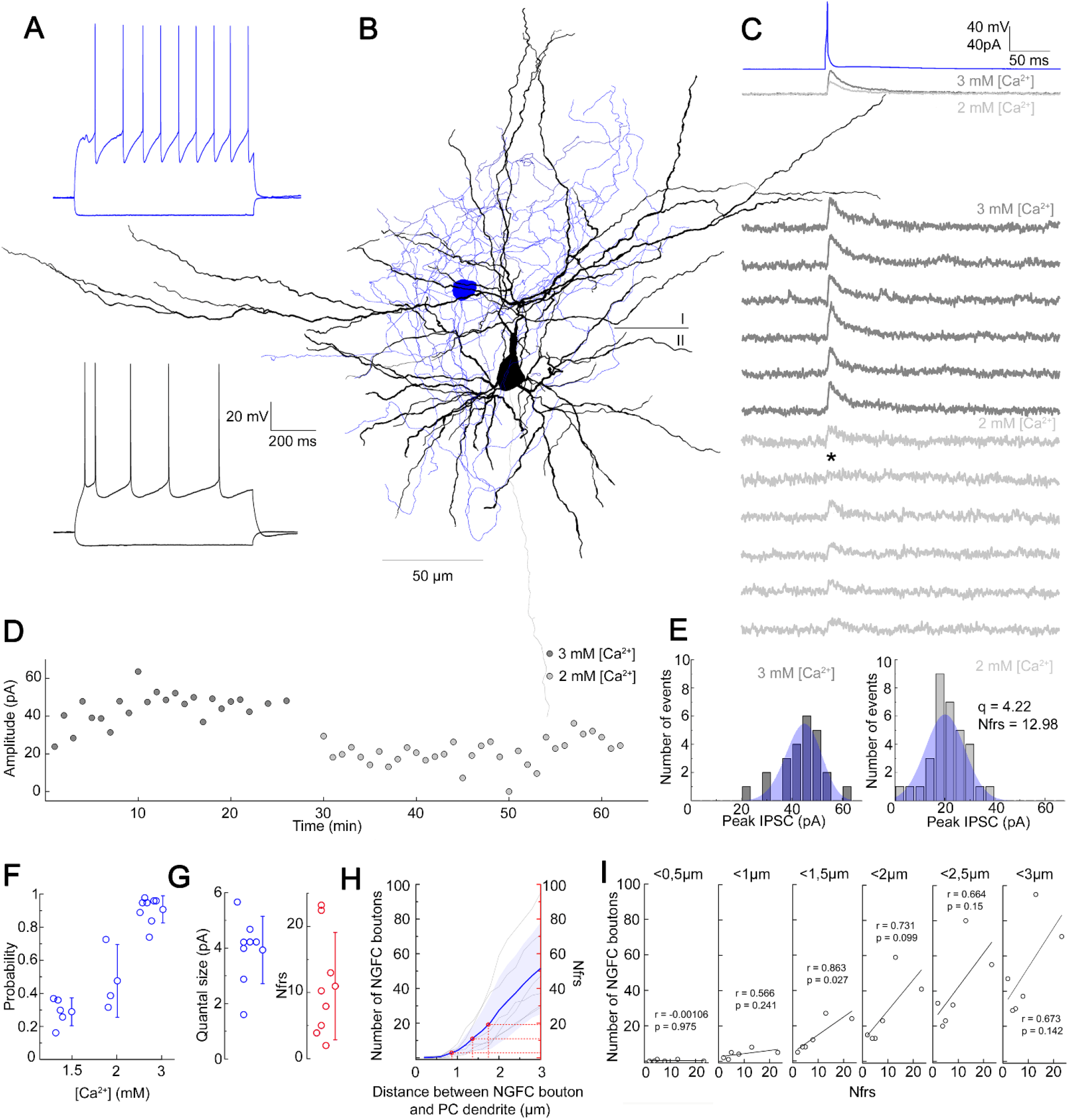
Quantal and structural characteristics of GABAergic connections established by individual neurogliaform cells. (A) Firing patterns of the presynaptic NGFC (blue) and postsynaptic PC (black). (B) Three-dimensional anatomical reconstruction of a recorded NGFC (soma and axon blue) and PC (soma and dendrites black, axon gray). (C) Presynaptic action potentials of the NGFC (top, blue) elicited of unitary IPSCs in the postsynaptic PC at −50 mV holding potential in different Ca^2+^ concentrations (middle, dark gray: 3 mM Ca^2+^, light gray 2 mM Ca^2+^). Bottom, representative consecutive traces of elicited unitary IPSCs. Asterisk marks synaptic transmission failure. (D) Single IPSC peak amplitudes recorded in high (3 mM Ca^2+^, dark gray) and low release probability conditions (2 mM Ca^2+^, light gray), respectively. (E) Distribution of IPSC peak amplitudes in 3 mM [Ca^2+^] (left) and 2 mM [Ca^2+^] (right), with projected binomial fits (blue). (F) Estimated release probability values in different experimental conditions (n= 8). (G) Estimated quantal size (3.93 ± 1.22 pA) and number of functional release sites (Nfrs; 10.96 ± 8.1) derived from Bayesian quantal analysis in each experiment (n= 8). (H) Number of NGFC boutons in the proximity of postsynaptic PC dendrites from anatomical reconstructions of connected NGFC to PC pairs (n= 8; gray, individual pairs; blue, average and SD). For comparison, red lines indicate mean ± SD of Nfrs shown on panel F corresponding to distances between presynaptic NGFC boutons and pyramidal cell dendrites. (I) Number of NGFC boutons counted at increasing distances from PC dendrites in NGFC to PC pairs. Correlation to Nfrs in the same pairs is best when counting boutons closer than 1.5 μm from PC dendrites.

### Structural characteristics of GABAergic connections established by the population of layer 1 neurogliaform cells

To have a better idea about how does the volume transmission radius potentially affect the fraction of converging outputs of L1 NGFC population to the same space, we developed a model to assess the overall output of NGFCs located in the supragranular layers of the neocortex. Unitary volume transmission by NGFCs is limited to their extremely dense axonal arborization (Oláh et al., 2009; Rózsa et al., 2017) Therefore, we determined the three-dimensional distribution of axon lengths of individual NGFCs with Sholl analysis (Fig. 2A). By superimposing individual NGFC reconstructions centered by their somata (n= 16; Fig. 2C) a representative distribution of axons was calculated as a function of distance from the soma Fig. 2D). We also assessed the distance between axonal boutons (n= 1456) along reconstructed axons of NGFCs (n= 6) and found that interbouton distances were 3.36 ± 2.54 μm (Supplementary Fig. 1). Next, we developed an algorithm that generates model NGFCs (n= 52) by growing axon arborizations similar (p= 0.99, two-sided K-S test, Fig. 2D) to the population of the experimentally reconstructed representative distribution of NGFC axons (n= 16) using interactions of segment lengths, branch point locations and segment orientations while keeping the density of axonal boutons along axon segments (Fig. 2B, C). In order to achieve a relatively complete representation of all NGFC axon terminals in a model at the populationlevel, we performed immunhistochemical labelling of α-actinin2, which is known to label the overwhelming majority of supragranular NGFCs in the neocortex (Uematsu et al., 2008). Somata immunoreactive for α-actinin2 in superficial cortical layers showed distribution along the axis perpendicular to the surface of the cortex with a peak at ~50-150 μm distance from the pia mater (Fig. 2E). According to this radial distribution and with no apparent tendency along the horizontal axis we placed NGFC somatas in a 354 x 354 x 140 μm volume to create a realistic spatial model of L1 NGFC population (Fig. 2F). Three dimensional pairwise shortest distances between α-actinin2+ somata (n= 152) and distances between somata placed into the model space (n= 374) were similar (p= 0.51, two-sided K-S test, Fig. 2G). We then used the axon growing algorithm detailed above from each soma position to model a population-wide distribution of NGFC axonal release sites. Quantal and structural properties of NGFC to PC connections shown above suggest a volume transmission distance of ~1.5 μm from potential sites of release (Fig. 1H, I), thus we mapped the coverage of surrounding tissue with GABA simultaneously originating from all NGFC terminals with a 1.5 μm of transmitter diffusion in the model. Using these conditions in simulations (n= 36), less than 8 NGFC axonal terminals contributed as effective GABA sources at any location in the superficial neocortex (Fig. 2H). Moreover, these boutons originated from a limited number of presynaptic NGFCs; when considering the extreme case of population-level cooperativity, i.e. when all putative NGFCs were active, most frequently a single NGFC release site serve as a GABA source (67.7 ± 7%) and potential interactions between two, three or more different NGFCs take place in limited occasions (15.34 ± 2.1%, 8.5 ± 2.6% and 8.45 ± 3.16%, respectively). The outcome of these simulations is consistent with earlier results suggesting that single cell driven volume transmission covers only the close proximity of NGFCs (Oláh et al., 2009) but also indicates potential interactions between a restricted number of neighbouring NGFCs.

**Figure 2.**
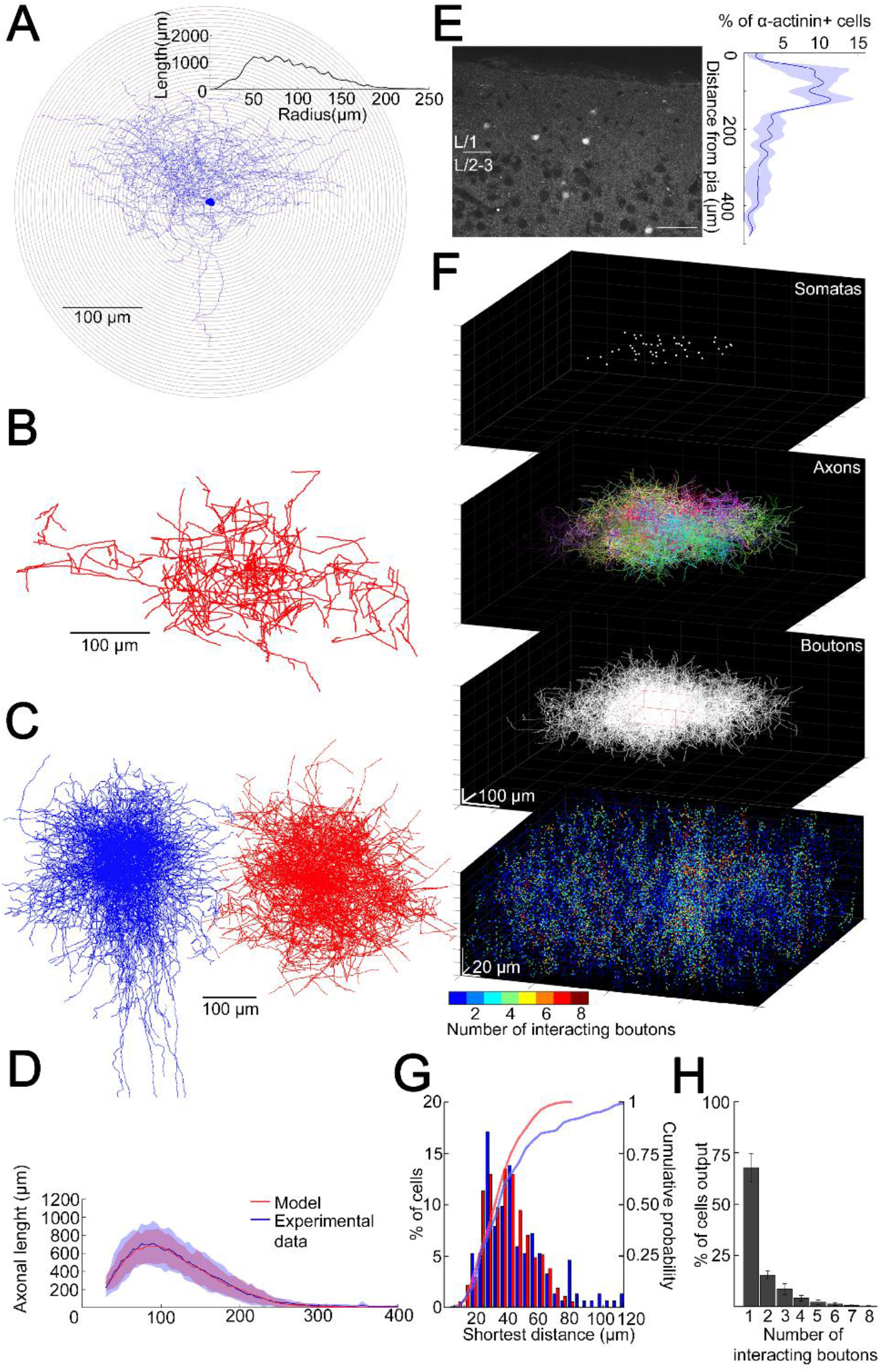
Structural characteristics of collective GABAergic output formed by the population of layer 1 neurogliaform cells. (A) Sholl analysis on the axonal arborization of an individual NGFC. Inset, axonal lengths measured in concentric shells of increasing radius (step, 10 μm). (B) Three-dimensional arborization of a model generated axon. (C) Superimposition of three-dimensionally reconstructed axonal arborizations of NGFCs (n= 16, blue) and the computer generated model NGFCs (n= 16, red) aligned at the center of somata. (D) Comparison of manually reconstructed axonal arborizations of NGFCs (n= 16; blue, mean; light blue, SD) and model generated axons (n= 52; red, mean; light red, SD) (E) Left, α-actinin2 immunohistochemistry in supragranular layers of the neocortex. Right, distribution of α-actinin2 immunopositive somata. (F) Top, three-dimensional model of NGFCs somata, axonal arborizations and bouton distributions in a 354 x 354 x 140 μm volume. Bottom, heat map showing the number of axonal boutons interacting at distances of < 1.5 μm. (G) Distribution of the shortest distance between somata in the model (red) and in α-actinin2 immunohistochemistry experiments (blue). (H) Percentage distribution of the number of interacting boutons within 1.5 μm distance from each NGFC.

### Coactivation of putative neurogliaform cells in L1 somatosensory cortex *in vivo*

Transcallosal fibers establish interhemispheric inhibition that operates via GABA_B_ receptor activation located on apical dendrites (L. M. Palmer et al., 2012) and it has been suggested that this massive GABA_B_ receptor recruitment in the superficial layers includes the activation of NGFCs (L. Palmer, Murayama, & Larkum, 2012). To assess the fraction of synchronously active putative NGFCs under close to physiological conditions, we applied *in vivo* two-photon Ca^2+^ imaging. We monitored the activity of L1 neurons bulk-loaded with calcium indicator Oregon Green BAPTA-1-AM (OGB-1-AM) (Fig. 3A) during hindlimb stimulation, which results in the activation of transcallosal inputs in L1 of the somatosensory cortex of urethane-anaesthetized rats (n= 6). Stimulation of the ipsilateral hindlimb (200 mA, 10ms) evoked Ca^2+^ signals in a subpopulation of neurons in L1 (n= 114 neurons; n= 46 vs. 68 responsive vs. non-responsive neurons, respectively; data pooled from six animals; Fig. 3B, 3C). On average 38.2 ± 5.2% of the L1 neurons were active following ipsilateral hindlimb stimulation, which is remarkably similar to the proportion found earlier (L. M. Palmer et al., 2012) (Fig. 3E). To further identify L1 neurons active during hindlimb stimulation, we performed immunohistochemistry for the actin-binding protein α-actinin2, (Uematsu et al., 2008) (Fig. 3F) using the same cortical area of L1 on which two-photon imaging was performed previously. Cross examination of neurons responsive/non-responsive to hindlimb stimulation versus neurons immunopositive/negative for α-actinin2 revealed that the majority of the active neurons were α-actinin2 positive (10 out of 15 neurons, 67%, n= 2 animals) and the majority of inactive neurons were α-actinin2 negative (22 out of 26 neurons, 85%, Fig. 3G) suggesting that a substantial fraction of L1 NGFCs are activated during hindlimb stimulation.

**Figure 3.**
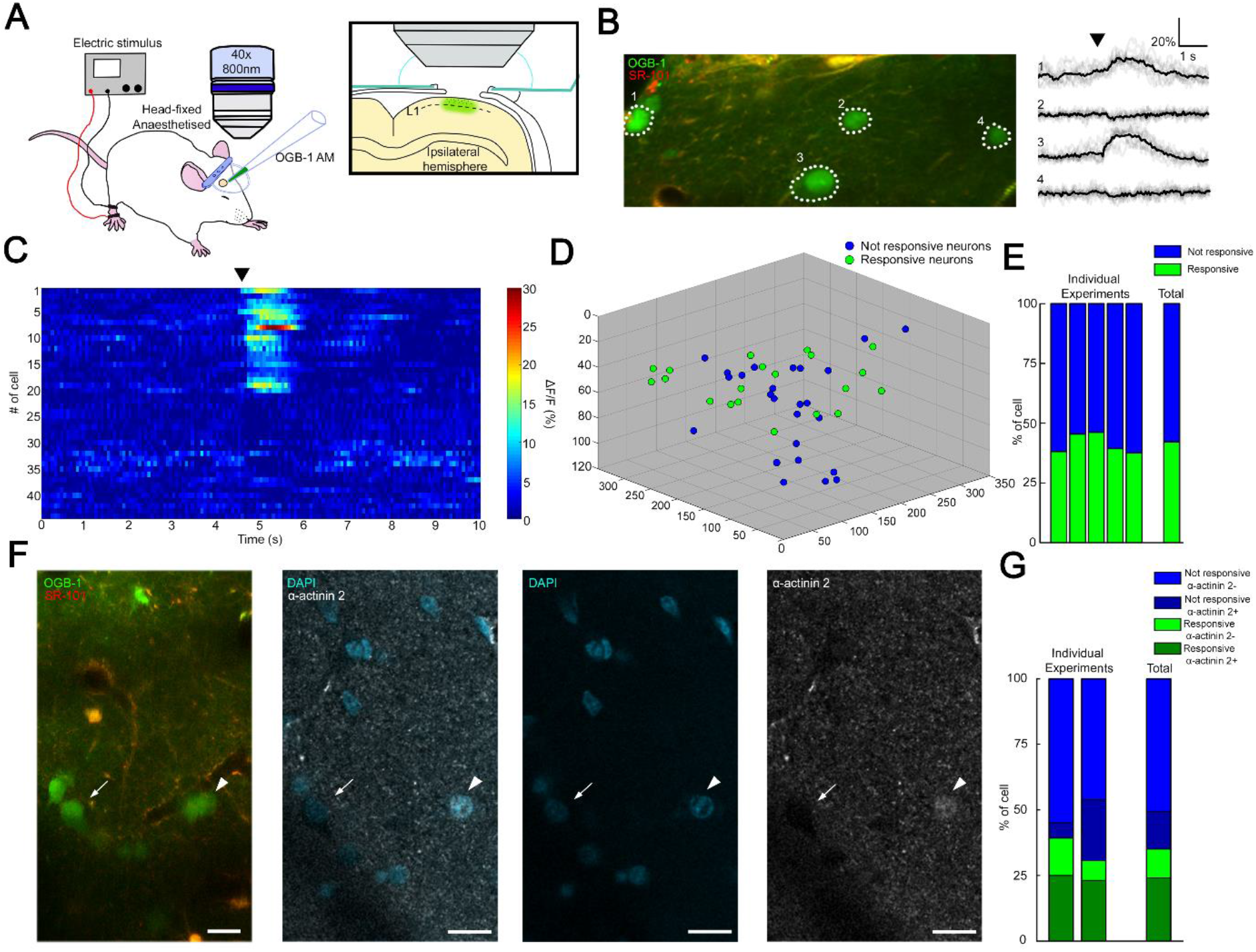
Coactivation of neurogliaform cell population in L1 somatosensory cortex *in vivo*. (A) Experimental setup. Head-fixed anaesthetized rats were placed under a two-photon microscope having a cranial window above the hindlimb somatosensory cortex. OGB-1 AM and SR 101 were injected into L1. Ipsilateral hindlimb stimulation was performed with an electric stimulator. (B) Two-photon image of neurons that were labeled with OGB-1 in L1. SR 101 labeled astrocytes. Right, ΔF/F changes of Ca^2+^ signals (grey: individual traces; black: mean of 10 consecutive traces) during series of ipsilateral stimulation (black arrowhead). Traces correspond to the marked cells. (C) Time-series heat map of 44 L1 interneurons evoked ΔF/F changes in Ca^2+^ signals during ipsilateral hindlimb stimulation (black arrowhead). (D) Scatter plot showing the spatial location of L1 interneuron somata. Colors are corresponding to the responsiveness (Not responsive, blue dots; responsive, green dots). (E) Stack columns show the fraction of responsive versus not responsive cells in different experiments (n= 5 animals). Far-right columns show the mean value. (F) *In vivo* two-photon image showing imaged neurons. To the right, confocal images from the same area shows immunohistochemical detection of α-actinin2+ neurons (arrowhead). α-actinin2-cells were visualized by exclusive DAPI staining (arrow). Scale bar, 20 μm. (G) Stack columns show the proportion of α-actinin2 immunoreactivity among responsive versus not responsive cells (n= 2 animals). Far-right columns show the mean value.

### Summation of convergent, unitary IPSPs elicited by NGFC

Our *in vivo* measurements above corroborate earlier results (L. M. Palmer et al., 2012) on widespread simultaneous activation of putative L1 NGFCs in response to transcallosal inputs. To directly measure the summation of converging inputs from superficial NGFCs, *in vitro* simultaneous triple recordings were performed from two presynaptic NGFCs and a target PC (n= 4, Fig. 4A). First we measured the amplitude of unitary IPSPs (n= 8) elicited by single NGFCs in the target PC and found that smaller and bigger inputs in a triplet were −1.68 ± 1.51 mV and −2.19 ± 1.33 mV, respectively. Next we activated the two NGFC inputs synchronously (0.17 ± 0.05 ms) and such coactivation resulted in moderately sublinear summation of convergent IPSPs (maximal nonlinearity, −9.1 ± 4.3 %) measured as the difference of calculated (−3.81 ± 2.76 mV) and experimentally determined (−3.57 ± 2.55 mV) sums of convergent single inputs (n= 4; Fig. 4B). These results are in line with experiments showing moderately sublinear interactions between identified, single cell evoked fast IPSPs (Tamás, Szabadics, & Somogyi, 2002). Interestingly, the time course of sublinearity followed the fast, presumably GABA_A_ receptor-mediated part of the unitary and summated IPSPs (Fig. 4C) suggesting that ionotropic and metabotropic GABAergic components of the same input combinations might follow different rules of summation. To test the interaction of unitary GABAB receptor-mediated IPSPs directly, we repeated the experiments above with the application of the GABA_A_ receptor antagonist gabazine (10 μM). Pharmacological experiments on the output of NGFCs are very challenging due to the extreme sensitivity of NGFC triggered IPSPs to presynaptic firing frequency (Capogna, 2011; Tamás, Simon Anna, & Szabadics, 2003) forcing us to collect the data in a different set of triple recordings (n= 8, Fig. 4D). As expected (Tamás et al., 2003) unitary, gabazine insensitive, slow IPSPs had onset latencies, rise times and half-widths similar to GABAB receptor-mediated responses (49.42 ± 5.8 ms, 86.95 ± 8.82 ms, and 252.27 ± 36.92 ms, respectively, n= 16, Fig. 4E). Peak amplitudes of converging smaller and bigger slow IPSPs were −0.66±0.22 mV and −0.94±0.37 mV respectively. Synchronous activation of two presynaptic NGFC converging onto the same pyramidal cell resulted in linear (−1.6 ± 6.6%) summation of slow IPSPs as peak amplitudes of calculated sums of individual inputs versus experimentally recorded compound responses were −1.58 ± 0.53 mV and −1.60 ± 0.55 mV, respectively (Fig. 4F). Taken together, our triple recordings in gabazine versus control conditions suggest linear interactions between slow, GABA_B_ IPSPs as opposed to sublinearly summating fast, GABA_A_ IPSPs elicited by the same presynaptic interneuron population.

**Figure 4.**
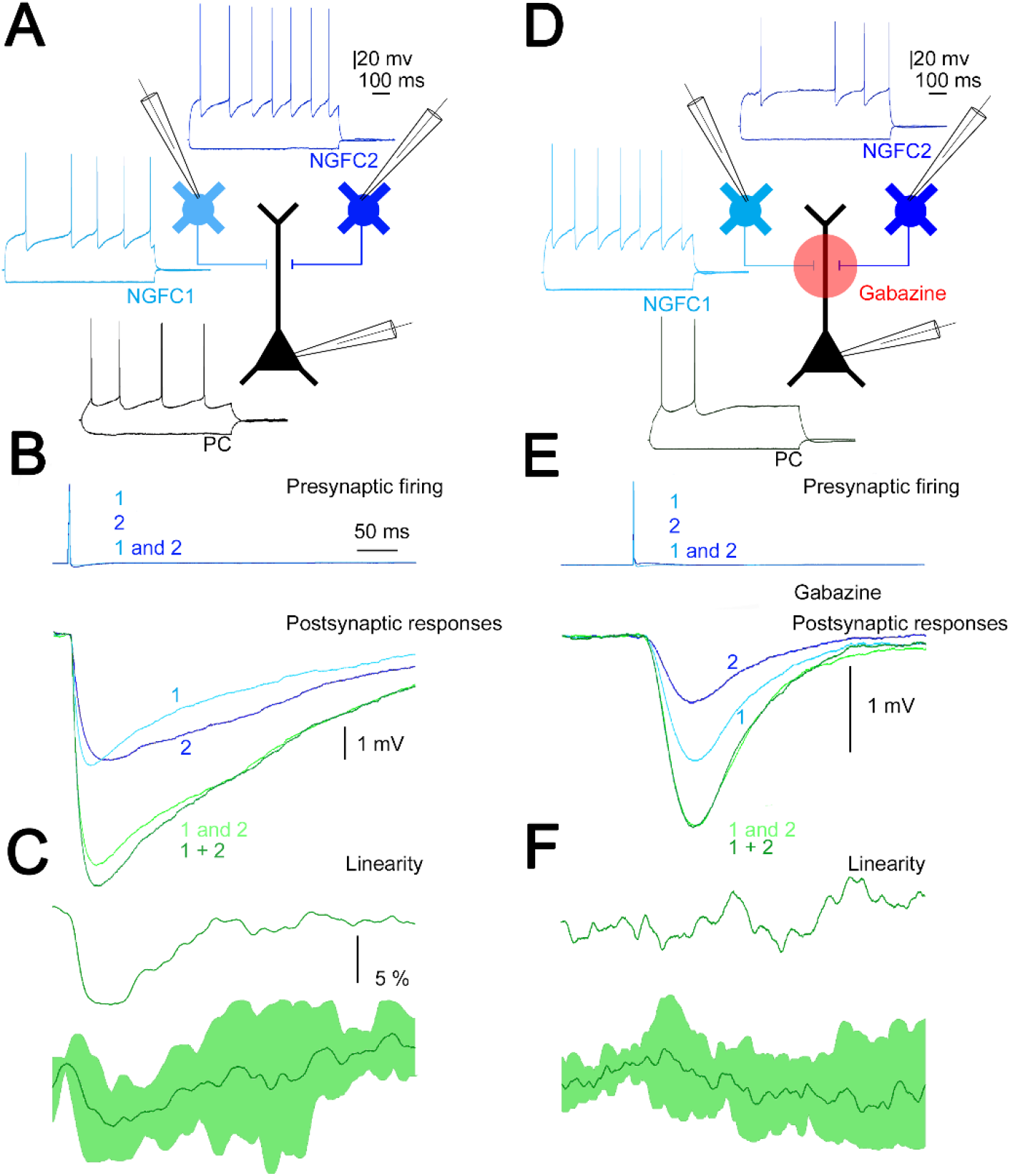
Summation of convergent, unitary IPSPs elicited by neurogliaform cells. (A) Schematic experimental setup of triplet recordings. Firing pattern of two presynaptic NGFCs (light blue and blue) and a postsynaptic pyramidal cell (black). (B) Action potential triggered under control conditions in the NGFCs individually (1, 2) or synchronously (1 and 2) elicited unitary (1, 2) and convergent (1 and 2) IPSPs in the postsynaptic PC. Below, the time course of the difference between the measured (1 and 2) and calculated (1 + 2) sums of convergent IPSPs. (C) The linearity of response summation on populations of convergent NGFC triggered IPSPs recorded in control conditions (n= 4) (dark green, population average; light green, SD). (D) Same as experimental setup as (A) but in the presence of GABA_A_ receptor antagonist, gabazine. (E) Identical stimulation protocol as (B), note the disappearance of the difference between the measured (1 and 2) and calculated (1 + 2) sums of convergent IPSPs. (F) Same as (C), but under blocking GABA_A_ receptors with gabazine (n= 8) (dark green, population average; light green, SD).

### Integration of GABA_B_ receptor-mediated responses are not affected by HCN channel and GABA reuptake

The predominant target area of the superficial NGFCs, the distal apical dendritic membrane of PCs, express voltage-dependent hyperpolarization-activated cyclic nucleotide-gated channel 1 (HCN1) known to attenuate dendritic signals (Berger, Larkum, & Lüscher, 2001; Kalmbach et al., 2018; Lörincz, Notomi, Tamás, Shigemoto, & Nusser, 2002; Robinson & Siegelbaum, 2003; Sheets et al., 2011). To investigate whether HCN1 channels contribute to mechanisms of interactions between GABA_B_ receptor-mediated postsynaptic responses we performed experiments on NGFC-to-PC pairs and evoked 1 to 4 action potentials (APs) in a single presynaptic NGFC at 100 Hz. This experimental configuration mimics the extreme conditions when multiple presynaptic release sites converge in a tight space and creating excessive GABAB receptor mediated inhibition. (Fig 5A). Triggering a single spike in the presence of gabazine (10 μM) did not saturate postsynaptic GABA_B_ receptors since the postsynaptic response induced by two spikes was proportional to the arithmetic sum of unitary postsynaptic responses (experimental sums: −1.25 mV calculated sums: −1.26 mV), apparently showing linear summation properties similar to triple recordings testing summation convergent inputs above. However, further increase in the number of evoked APs to 3 and 4 introduced sublinearity to summation (n= 6, 1AP: −0.63 ± 0.50 mV, 2APs: −1.25 ± 1.06 mV, 3APs: −1.53 ± 0.84 mV, 4APs: −1.61 ± 1.09 mV, Fig. 5B; normalized values: 2APs: 2.00 ± 1.08, 3APs: 2.34 ± 1.16, 4APs: 3.17 ± 1.26, Fig. 5C). Importantly, recordings in the presence of HCN1 channel blocker, ZD-2788 (10 μm) showed summation properties similar to control, summation was linear with two APs and changed to slightly sublinear upon the 3rd to 4th spike, (n= 5, 1AP: −0.82 ± 0.63 mV, 2APs: −1.59 ± 0.76 mV, 3APs: −1.66 ± 0.72 mV, 4APs: −1.90 ± 1.07 mV, Fig. 5B; normalized values and its comparison to control: 2APs: 2.06 ± 1.06, p= 0.983; 3APs: 1.99 ± 1.17, p= 0.362; 4APs: 2.56 ± 1.6, p= 0.336; two-sided MW U test, Fig. 5C). These experiments suggest that when a physiologically probable number of NGFCs are simultaneously active, HCN1 channels are locally not recruited to interfere with the summation of GABA_B_ receptor-mediated responses.

**Figure 5.**
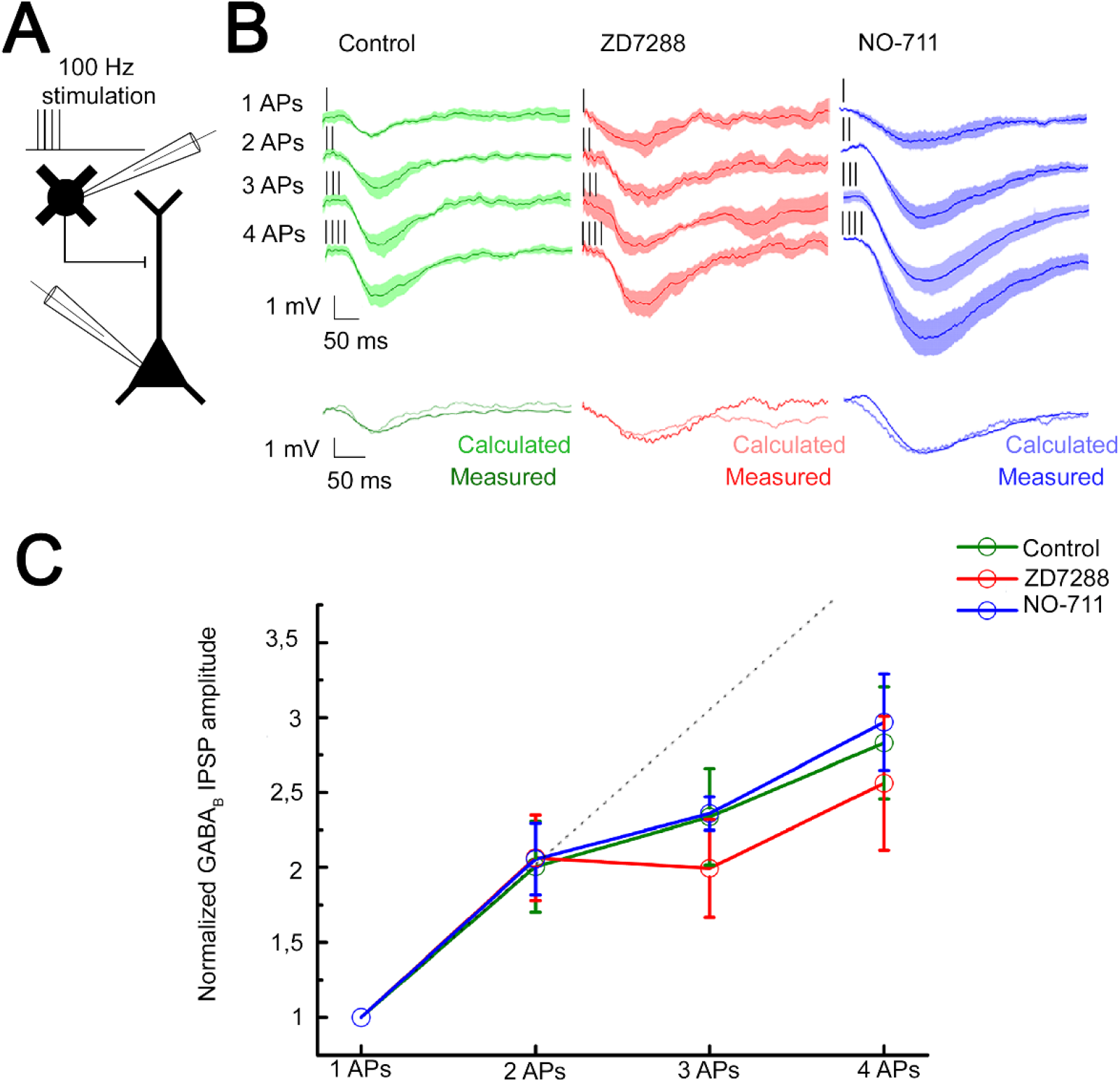
Integration of GABA_B_ receptor-mediated responses are not affected by HCN channel and GABA reuptake. (A) Schematic experimental setup of paired recordings. Bursts of up to four action potentials (APs) were elicited in NGFCs at 100 Hz in the presence of gabazine. (B) NGFC to PC paired recordings showed similar linear GABA_B_ receptor-mediated summation under control conditions. Top, individual traces showing IPSP kinetics upon AP burst protocol (vertical lines indicating the triggered APs) during control (green traces, n= 6), in presence of hyperpolarization-activated cation (HCN) channel blocker ZD7288 (red traces, n= 5) or GABA reuptake blocker NO-711 (blue traces, n= 6). Bottom, traces show measured IPSP from two consecutive presynaptic stimulation (measured) and the arithmetic sum of two unitary IPSP (calculated). (C) Summary of normalized IPSP peak amplitudes. Compare to control conditions (2APs: 2.00 ± 1.08; 3APs: 2.34 ± 1.16; 4APs: 3.17 ± 1.26) summation properties of GABA_B_ mediated unitary IPSPs are not effected by application of ZD7288 (2APs: 2.06 ± 1.06, p= 0.983; 3APs: 1.99 ± 1.17, p= 0.362; 4APs: 2.56 ± 1.6, p= 0.336; two-sided MW U test) neither NO-711 (2APs: 2.06 ± 1.17, p= 0.853; 3APs: 2.36 ± 0.31, p= 0.645; 4APs: 2.9 7± 1.54, p= 0.515; two-sided MW U test). Dashed line indicates the linearity.

Previous experiments suggested that a single AP in a NGFC is able to fill the surrounding extracellular space with an effective concentration of GABA (Oláh et al., 2009) and, in turn, extracellular GABA concentration producing GABA_B_ receptor activation is tightly regulated via GABA transporters (GAT-1) (Gonzalez-Burgos, Rotaru, Zaitsev, Povysheva, & Lewis, 2009; Isaacson et al., 1993; Rózsa et al., 2017; Szabadics, Tamas, & Soltesz, 2007). Therefore, we tested whether GAT-1 activity affects the summation of GABA_B_ receptor-mediated responses potentially limiting the number of GABA_B_ receptors reached by GABA released by NGFCs. Selective blockade of GAT-1 with NO-711 (10 μm) increased the amplitude of GABA_B_ receptor-mediated IPSP, however, it did not influence summation properties (n= 6, 1AP: −1.11 ± 0.62 mV, 2APs: −2.28 ± 1.07 mV, 3APs: −3.1 ± 0.40 mV, 4APs: −3.54 ± 1.59 mV, Fig. 5B; normalized values and its comparison to control: 2APs: 2.06 ± 1.17, p= 0.853; 3APs: 2.36 ± 0.31, p= 0.645; 4APs: 2.97 ± 1.54, p= 0.515; two-sided MW U test, Fig. 5C). Accordingly, interactions between an *in vivo* realistic number of simultaneously active NGFCs lead to linear GABA_B_ response summation even if increased concentration of GABA is present in the extracellular space.

### Subcellular localization of GABA_B_ receptor-GIRK channel complex determines summation properties

High resolution quantitative electron microscopy showed that GABA_B_ receptors and GIRK channels are segregated on dendritic shafts, however, receptor-channel complexes colocalize on dendritic spines (Á. Kulik et al., 2006). Theoretical studies suggest that the distance between the receptor and effector limits the recruitment of effector molecules to the vicinity of receptors (Brinkerhoff, Choi, & Linderman, 2008; Á. Kulik et al., 2006), thus we asked if linear summation was potentially a result of the locally constrained GABAB receptor - GIRK channel interaction when several presynaptic inputs converge. To this end, we constructed a simulation environment based on a previously published three-dimensional reconstruction of a postsynaptic dendritic segment (Edwards et al., 2014) targeted by realistically positioned release sites of NGFCs (Fig. 6A). Molecular interactions in this spatially realistic system were modeled using Monte Carlo algorithms to simulate movements and reactions of molecules (Kerr et al., 2008). Membranes of the postsynaptic dendritic segment were populated (see methods) with GABA_B_ receptors and GIRK channels according to compartment-dependent data from SDS-digested freeze-fracture replica immunolabeling (Á. Kulik et al., 2006) (Fig. 6C). Neurotransmitter diffusion in the brain is influenced by tissue tortuosity and the fraction of extracellular space in total tissue volume (Sykova & Nicholson, 2008), thus we simulated realistic molecular diffusion in tortuous extracellular space (Tao, Tao, & Nicholson, 2005) (see methods). The number and position of NGFC presynaptic boutons around the postsynaptic dendritic segments in the model were used according to structural characteristics of GABAergic connections established by individual NGFCs (n= 4 boutons 1.2 ± 0.7 μm from the dendrite; Fig. 1I, H) and according to the bouton density determined for the overall output of NGFC population (Fig. 2F). Previous work suggests that a single AP in a NGFC generates GABA concentrations of 1 to 60 μM lasting for 20-200 ms (Karayannis et al., 2010) so we used a similar ~1 to 60 μM of GABA concentration range at 0.5 to 2 μm distance from the release sites (Supplementary Fig. 4) and GABA exposure times of 114.87 ± 2.1 ms with decay time constants of 11.52 ± 0.14 ms. Our modeling trials show that single AP triggered GABA release can activate a total of 5.82 ± 2.43 GABA_B_ receptors (2.81 ± 1.55 on spine, 3.01 ± 1.71 on the shaft). Furthermore, activation of GABA_B_ receptors triggers intracellular mechanisms and the initial GDP/GTP exchange at the Gα subunit separates the G-protein heterotrimeric protein and produces Gβγ subunits (peak number of Gβγ subunits for single AP: 338.54 ± 138.75). Lateral membrane diffusion of Gβγ subunits lead to the activation of 3.66 ± 2.17 GIRK channels in total (2.47 ± 1.88 on spine, 1.17 ± 1.26 on the shaft) in response to single AP. Next, consecutive GABA releases were induced with 10 ms delays to replicate the 100 Hz stimulation protocol used in the experiments above (Fig. 5A). The increased GABA concentration from two sequential stimuli raised the number of active GABA_B_ receptors to 11.29 ± 3.48 (5.57 ± 2.36 on spine, 5.72 ± 2.52 on the shaft). Three and four consecutive releases activated a total of 16.19 ± 3.88 and 20.99 ± 4.99 GABA_B_ receptors, respectively (7.96 ± 2.74 on spine, 8.23 ± 2.97 on the shaft and 10.62 ± 3.28 on spine, 10.37 ± 3.53 on the shaft, respectively). When modeling consecutive GABA releases, massive amount of Gβγ subunits were produced together with a decline in relative production efficacy per APs, possibly due to the limited number of G-proteins serving as a substrate in the vicinity of active receptor clusters (peak number of Gβγ subunits for 2AP: 612.10 ± 171.95; 3AP: 857.78±194.14; 4AP: 1081.81 ± 229.57). Two consecutive APs resulted in the activation of 6.98 ± 3.29 GIRK channels (2.13 ± 1.63 on dendritic shaft and 4.85 ± 2.65 on spine) in the simulations. Importantly, this number of activated GIRK channels in response to two APs was close to the arithmetic sum of the number of GIRK channels activated by two single AP responses (−4.87% in total, −1.86% on spines and −9.86% on the shaft; Fig. 4E, 5A). Further increase in the GABA exposure proportional to three and four action potentials lead to the activation of 10.39 and 12.89 GIRK channels, respectively (7.01 ± 3.11 and 8.68 ± 3.46 on spines and 3.37 ± 2.08 and 4.21 ±2. 48 on the shaft, respectively). These numbers of GIRK channels corresponded to −5.68 and −13.58% of the arithmetic sum of GIRK channels activated by three and four single AP responses (−5.71 and 13.82% on spines and −4.15 and −11.16% on the shaft).

**Figure 6.**
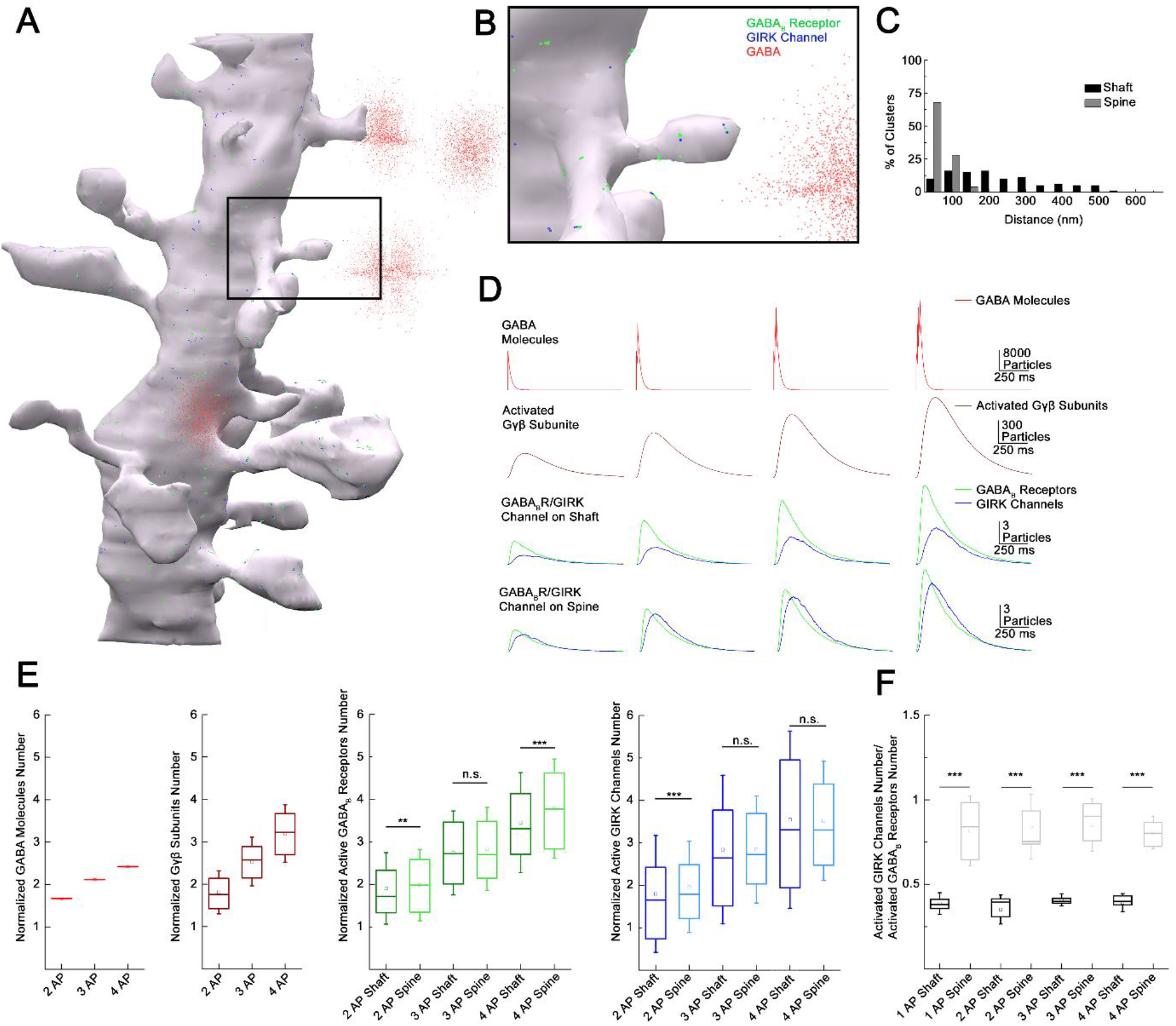
Subcellular localization of GABA_B_ receptor-GIRK channel complex determines summation properties. (A) Visualization of the complete MCell based model in the course of GABA release. (B) Magnified view of the model. (C) Distribution of GABA_B_ receptors and GIRK channel clusters on the dendritic membrane in the model (grey bars: dendritic spine; black bars: dendritic shaft). (D) Overview of the molecular interactions during increasing GABA release. Top to bottom: NGFC output simulated by releasing GABA (red) in the extracellular space proportional to 1-4 action potential stimulation. Below, the total number of produced Gβγ subunits (brown) by activated GABA_B_ receptors (green) located on the dendritic shaft and spine. After lateral diffusion in the plasma membrane, Gβγ subunits bind to GIRK channels (blue). (E) Boxplot of GABA, Gβγ subunits, GABA_B_ receptors and GIRK channels quantity normalized to 1AP (GABA: 2APs: 1.67 ± 0.004, 3APs: 2.12 ± 0.005, 4APs: 2.42 ± 0.006, Gβγ subunits: 2APs: 1.85 ± 0.51, 3APs: 2.53 ± 0.57, 4APs: 3.2 ± 0.68; GABA_B_ receptor shaft: 2APs: 1.91 ± 0.84, 3APs: 2.74 ± 0.99, 4APs: 3.45 ± 1.18; GABAB receptor spine: 2APs: 1.98 ± 0.84, 3APs: 2.84 ± 0.98, 4APs: 3.78 ± 1.17; GIRK channel shaft: 2APs: 1.8 ± 1.37, 3APs: 2.84 ± 1.75, 4AP: 3.55 ± 2.09; GIRK channel spine: 2APs: 1.96±1.07, 3APs: 2.84 ± 1.26, 4APs: 3.52 ± 1.4). Square indicate the mean, line shows the median inside the boxplot. (F) Quantification of the signaling effectiveness on the shaft and spine region of the model dendrite during increasing GABA release (1AP: shaft: 0.39 ± 0.06, spine: 0.82 ± 0.21, p < 0.005, n= 1164, two-sided MW U test; 2AP: shaft: 0.35 ± 0.086, spine: 0.84 ± 0.19, p < 0.005, n= 534, two-sided MW U test; 3AP: shaft: ± 0.41 ± 0.036, spine: 0.85 ± 0.15, p < 0.005, n= 1871, two-sided MW U test; 4AP: shaft: 0.39 ± 0.05, spine: 0.81 ± 0.01, p < 0.005, n= 709, two-sided MW U test). Square indicate the mean, line shows the median inside the boxplot.

GABA_B_ receptor and GIRK channel complexes located in particular subcellular compartments appeared to have different effectiveness of recruiting GABA_B_ receptors and active GIRK channels in our simulations (Fig. 6D). We observed different numbers of GABA_B_ receptors activated on the shaft and spine (Normalized values to 1AP: 2APs: shaft: 1.91 ± 0.84, spine: 1.98 ± 0.84, n= 534, p= 0.009; 3APs: shaft: 2.74 ± 0.99, spine: 2.84 ± 0.98, n= 1871, p= 0.173; 3APs: shaft: 3.45 ± 1.18, spine: 3.78 ± 1.17, n= 709, p< 0.005, two-sided MW U test, Fig. 6E). The recruitment of GIRK channels was more effective on spines compared to shafts when triggering 2 APs (Normalized values to 1AP: shaft: 1.8±1.37, spine: 1.96±1.07, n=534, p<0.005); the trend was similar in response to three and four APs, but results were not significant (Normalized values to 1AP: 3AP: shaft: 2.84±1.75, spine: 2.84 ± 1.26, n= 1871, p= 0.109; 4AP: shaft: 3.55 ± 2.09, spine: 3.52 ± 1.4, n= 709, p= 0.216, two-sided MW U test, Fig. 6E). The compartment-specific effectiveness of signaling as the ratio of activated GIRK channels and active GABA_B_ receptors (Fig. 6F) shows that spines represent the preferred site of action corroborating earlier suggestions (Qian & Sejnowski, 1990).

## Discussion

The unique inhibitory communication via volume transmission separates NGFC interneurons from other interneuron classes in the neocortex. Numerous observations support the idea of volume transmission (Overstreet-Wadiche & McBain, 2015). (1) NGFC activation generates an unusually prolonged inhibition on the postsynaptic cell (Karayannis et al., 2010; Mańko, Bienvenu, Dalezios, & Capogna, 2012; Oláh et al., 2009; Szabadics et al., 2007). (2) Released GABA acts on synaptic and extrasynaptic GABA receptors (Karayannis et al., 2010; Oláh et al., 2009; Price, 2005; Tamas, 2003), (3) as well as on nearby presynaptic terminals (Oláh et al., 2009). (4) NGFCs show a very high rate of functional coupling between the neighboring neurons (Jiang et al., 2015; Oláh et al., 2009). (5) Ultrastructural observations showed the lack of clearly defined postsynaptic elements in the apposition of the NGFC boutons (Mańko et al., 2012; Oláh et al., 2009; Vida, Halasy, Szinyei, Somogyi, & Buhl, 1998). (6) NGFCs act on astrocytes within the reach of their axonal arborization through nonsynaptic coupling (Rózsa et al., 2017). The distance of effective operation through NGFC driven volume transmission, however, is not clear. Here, we used functional and structural characterization of NGFC-PC inhibitory connections and suggest that GABA released from NGFC axonal terminals activates GABA receptors up to about ~1.8 μm, a result remarkably similar to previous estimations for the range of extrasynaptic action of synaptically released GABA (Farrant & Nusser, 2005; Overstreet-Wadiche & McBain, 2015; Overstreet et al., 2000) Our experiments also shed light to some quantal properties of NGFC’s GABA release. These experiments are constrained by the robust use-dependent depression mediated by NGFCs (Karayannis et al., 2010; Tamás et al., 2003), therefore implementation of multiple probability fluctuation analysis (MPFA) (Silver, 2003), the gold standard for quantal analysis, was not feasible and BQA (Bhumbra & Beato, 2013b) was needed as an alternative. The revealed linear correlation between BQA-derived Nfrs and the number of NGFC boutons putatively involved in transmission is compatible with the release of a single docked vesicle from individual NGFC boutons and suggest that multivesicular release is not essential for GABAergic volume transmission.

The functional distance of volume transmission is particularly important for the characterization of interactions between NGFCs and for understanding the population output of NGFCs. Realistic representation of an entire subpopulation of neurons is considered essential for the interpretation of network functions (Karnani, Agetsuma, & Yuste, 2014; Markram et al., 2015) and pioneering full-scale data-driven models were effective in deciphering emerging functions of interneuron populations (Bezaire, Raikov, Burk, Vyas, & Soltesz, 2016). However, network diagrams addressing the function of NGFCs exclusively based on synaptic connectivity underestimate the spread of output without incorporating volume transmission by an order of magnitude (Oláh et al., 2009). Although the concept of blanket inhibition has been suggested for networks of interneurons populations having overlapping axonal arborizations and dense synaptic output (Karnani et al., 2014), our spatial model based on high resolution reconstructions of labeled NGFCs takes the concept to its extremes and reveals an unprecedented density of release sites for a population of cortical neuron and shows that the overwhelming majority of the superficial cortical space is effectively covered by at least one NGFC. At the same time, the redundancy of the NGFC population is limited and a single cortical spatial voxel is reached by GABA released from a limited number of individual NGFCs, ~83 % of space is covered by 1 or 2 NGFCs. Our relatively simple *in vivo* approach to gauge potential synchronous action of NGFCs gave positive results. This is in line with earlier observations suggesting widespread action of putative NGFCs in terminating persistent activity (Craig et al., 2013), or powerfully suppressing dendritic Ca^2+^ dynamics in L2/3 and L/5 (L. M. Palmer et al., 2012; Wozny & Williams, 2011). Strong cholinergic neuromodulation of NGFCs (Poorthuis et al., 2018) and frequent gap junctional coupling between NGFCs (Simon, Oláh, Molnár, Szabadics, & Tamás, 2005) further facilitates concerted action and are likely to play a major role in synchronizing the NGFC network (Yao et al., 2016).

When studying simultaneous action of NGFCs, our direct measurements of two converging NGFC inputs on L2/3 PC from simultaneous triple whole cell patch clamp recordings revealed sublinear summation properties for ionotropic GABA_A_ receptor mediated responses. These results support classic theories on synaptic input interactions (Jadi et al., 2012; Koch et al., 1983; London & Häusser, 2005; Qian & Sejnowski, 1990; Silver, 2010) and are in line with earlier experiments measuring interactions of anatomically identified inputs converging to neighboring areas of the dendritic tree of the same postsynaptic cell (Hao, Wang, Dan, Poo, & Zhang, 2009; Tamás et al., 2002). Mechanisms of interaction between convergent inputs from NGFCs might be similar to those suggested for short-term synaptic depression of GABA_A_ responses such as local drops in Cl^-^ driving force and membrane conductance (Huguenard & Alger, 1986; McCarren & Alger, 1985; Staley & Proctor, 1999). To our knowledge, the simultaneous triple recordings of two presynaptic NGFCs targeting the same postsynaptic PC represent the first direct experimental attempt addressing the summation of metabotropic receptor mediated postsynaptic interactions. To date, scarce computational model studies were aimed to explore the integration properties of GABA_B_ receptor-mediated responses and suggested a highly supralinear interaction through the amplification effect of G-protein cooperativity (Destexhe, 1995). Our experimental approach indicates linear interactions between GABA_B_ receptor mediated responses in case the number of converging presynaptic cells corresponds to the number of NGFCs cooperating during *in vivo* network operations. This suggests that converging afferents that act on inhibitory metabotropic receptors in the same postsynaptic voxel show linear or slightly sublinear summation, conserving the impact of individual inputs. However, we cannot exclude the possibility that widespread synchronization across various interneuron populations might shift the summation arithmetic in a nonlinear fashion.

Intrinsic properties of postsynaptic pyramidal cells might also contribute to the regulation of summation. HCN1 channels are known to be enriched in the distal dendrites of pyramidal cells and mediate K^+^ cationic current activated by membrane hyperpolarization (Kalmbach et al., 2018; Lörincz et al., 2002; Robinson & Siegelbaum, 2003) Our experiments presented above show that summation properties in response to synchronized inputs from NGFCs are not significantly affected by HCN1 channels, presumably due to the relatively moderate local hyperpolarization arriving from NGFCs; again, further studies are needed to test the influence of additional interneuron classes coactivated together with NGFCs. We predict that further GABAergic activity is unlikely to change summation arithmetics based on our negative results when blocking the high-affinity plasma membrane GABA transporters concentrated in the perisynaptic and extrasynaptic areas (Melone, Ciappelloni, & Conti, 2015) effective in modulating GABA-mediated inhibition trough extrasynaptic GABA spillover (Barbour & Häusser, 1997; Hamann, Rossi, & Attwell, 2002; Scanziani, 2000; Szabadics et al., 2007). Despite having similar summation arithmetics of two consecutive APs to the triple-recording configuration, it remains undefined as to what extent multiple presynaptic APs resemble synchronous activation of individual release sites. Presynaptic GABA_B_ receptor-mediated decrease in Ca^2+^ is unlikely (Karayannis et al., 2010), however, depletion of the readily releasable pool of vesicles leading to synaptic depression cannot be ruled out. As suggested by pioneering simulations on the summation of GABA_B_ receptor-mediated signaling (Destexhe, 1995), a crucial intrinsic factor in the postsynaptic cells is the molecular cascade linking GABA_B_ receptors to the GIRK channels through G-proteins. Our experimental evidence for close to linear or slightly sublinear summation of GABA_B_ receptor-mediated responses suggests that even if amplification through G-proteins plays a role, it is unable to overturn local membrane or K^+^ concentration dependent factors promoting sublinearity (Dascal, 1997; Inanobe & Kurachi, 2014; Stanfield et al., 2002; Wickman & Clapham, 1995). Amplification of GIRK current by G-proteins could be hampered by the need of cooperative action of up to four G-protein βγ subunits to be effective in opening GIRK channels. In addition, hyperpolarization and the accompanying relatively low [Na^+^]i might also limit GIRK channel activation knowing that high [Na^+^]i promotes GIRK channel opening in depolarized cells (Wang et al., 2016). The latter scenario might promote a brain state-dependent summation of metabotropic inhibitory signals in active neuronal networks, that remains to be tested in future experiments. On the other hand, our ultrastructural model corroborates pioneering suggestions (Á. Kulik et al., 2006; Qian & Sejnowski, 1990) that the effect of GABA_B_ receptors is more prominent on dendritic spines compared to dendritic shafts, having approximately twice the number of activated GIRK channels per GABA_B_ receptor on spines versus shafts. Admittedly, our simulations could not cover the extensive intracellular signaling pathways known to be influenced by GABA_B_ receptors (Gassmann & Bettler, 2012; Padgett & Slesinger, 2010; Terunuma, 2018) and future availability of comprehensive transporter and extracellular space distributions of layer 1 would enrich the model (Hrabetova, Cognet, Rusakov, & Nägerl, 2018; Korogod, Petersen, & Knott, 2015; Pallotto, Watkins, Fubara, Singer, & Briggman, 2015). Nevertheless, our experiments and simulations suggest that nonsynaptic GABAergic volume transmission providing relatively homogeneous and sufficient concentrations of GABA combined with increased clustering of GABA_B_ receptors and on spines compared to shafts governs compartment dependent efficacy.

Taken together, our experimental results and modeling analysis suggest that a randomly chosen location in the neuropil of layer 1 is targeted by a moderate number (usually one or two) presynaptic NGFCs. In turn, there is no apparent gap in the neurogliaform coverage of layer 1, i.e. most elements of the neuropil including classic postsynaptic compartments, presynaptic terminals or non-neuronal cells are located sufficiently close to terminals of at least one NGFC and receive GABA nonsynaptically. Interestingly, when two NGFCs which share target territory are coactivated or a single NGFC has a limited number of consecutive spikes, linear arithmetics accompany GABA_B_ receptor summation. This supports the hypothesis that the density and distribution of neocortical NGFCs and their axonal terminals combined with the effective range of GABAergic volume transmission appear optimized for a spatially ubiquitous and predominantly linear metabotropic GABA_B_ receptor summation.

## Acknowledgement

This work was supported by the ERC INTERIMPACT project, by the Hungarian Academy of Sciences, the Hungarian National Office for Research and Technology GINOP 2.3.2-15-2016-00018, Élvonal KKP 133807 and the National Brain Research Program, Hungary. We would like to thank Dr. Angus Silver for the useful comment on an early version of the manuscript, Dr. Chandrajit Bajaj for the permission of using the 3D reconstruction of a dendritic structure and Éva Tóth and Nelli Ábrahám Tóth for their exceptional technical assistance.

## Author Contributions

Conceptualization, G.T.; Methodology, A.O.,G.K.,G.M. and G.T.; Investigation, A.O.,G.K.,G.O. and J.B.; Software, A.O.; Formal Analysis, A.O., G.K., G.O. and G.T.; Writing – Original Draft, A.O., G.K. and G.T.; Writing – Review & Editing, G.T.; Visualization, A.O.,G.K. and G.T.; Funding Acquisition, G.T.; Supervision, G.T.

## Methods

### Slice preparation

Experiments were conducted to the guidelines of University of Szeged Animal Care and Use Committee. We used young adult (19 to 46-days-old, (P) 23.9 ± 4.9) male Wistar rats for the electrophysiological experiments. Animals were anaesthetized by inhalation of halothane, and following decapitation, 320 μm thick coronal slices were prepared from the somatosensory cortex with a vibration blade microtome (Microm HM 650 V; Microm International GmbH, Walldorf, Germany). Slices were cut in ice-cold (4°C) cutting solution (in mM) 75 sucrose, 84 NaCl, 2.5 KCl, 1 NaH_2_PO_4_, 25 NaHCO_3_, 0.5 CaCl_2_, 4 MgSO_4_, 25 d (+)-glucose, saturated with 95% O_2_ and 5% CO_2_. The slices were incubated in 36°C for 30 minutes, subsequently the solution was changed to (in mM) 130 NaCl, 3.5 KCl, 1 NaH_2_PO_4_, 24 NaHCO_3_, 1 CaCl_2_, 3 MgSO_4_, 10 d (+)-glucose, saturated with 95% O_2_ and 5% CO_2_, and the slices were kept in it until experimental use. The solution used for recordings had the same composition except that the concentrations of CaCl2 and MgSO_4_ were 3 mM and 1.5 mM unless it is indicated otherwise. The micropipettes (3-5 MΩ) were filled (in mM) 126 K-gluconate, 4 KCl, 4 ATP-Mg, 0.3 GTP-Na2, 10 HEPES, 10 phosphocreatine, and 8 biocytin (pH 7.25; 300 mOsm).

### In vitro Electrophysiology and Pharmacology

Somatic whole-cell recordings were obtained at ~37°C from simultaneously recorded triplets and doublets of NGF and PC cell visualized by infrared differential interference contrast video microscopy at depths 60-160 μm from the surface of the slice (Zeiss Axio Examiner LSM7; Carl Zeiss AG, Oberkochen, Germany), 40x water-immersion objective (1.0 NA; Carl Zeiss AG, Oberkochen, Germany) equipped with Luigs and Neumann Junior micromanipulators (Luigs and Neumann, Ratingen, Germany) and HEKA EPC 10 patch clamp amplifier (HEKA Elektronik GmbH, Lambrecht, Germany). Signals were filtered 5 kHz, digitalized at 15 kHz, and analyzed with Patchmaster software.

Presynaptic cells were stimulated with a brief suprathreshold current pulse (800 pA, 2-3 ms), derived in >60 s interval. In experiments, where two presynaptic NGFC were stimulated simultaneously the interval was increased >300 s. The stimulation sequence, in which one or the other or both presynaptic NGFCs were stimulated was constantly altered, therefore the potential rundown effect or long term potentiation would affect all three stimulation condition equally. In the case of 100 Hz presynaptic burst stimulation the interval was increased >300 s. During stimulation protocol, the order of triggering a set of 1 to 4 APs on the NGFc were randomized. The postsynaptic responses were normalized to the single AP in each individual set. During postsynaptic current-clamp recording −50 mV holding current was set. The experiments were stopped if the series resistance (Rs) exceeded 35 MOhm or changed more than 20%. During postsynaptic voltage-clamp recordings, Rs and whole-cell capacitance were monitored continuously. The experiment was discarded if the compensated Rs change reached 20% during recording.

Pharmacological experiments were carried out on NGFC-PC pairs using ACSF with the following drugs: 10 μM SR 95531 hydrobromide (Tocris), 10 μM D-(−)-2-Amino-5-phosphonopentanoic acid (D-AP5) (Tocris), 10 μM 2,3-Dioxo-6-nitro-1,2,3,4-tetrahydrobenzo[*f*]quinoxaline-7-sulfonamide (NBQX) (Tocris), 10 μM 4-(N-Ethyl-N-phenylamino)-1,2 dimethyl-6-(methylamino) pyrimidinium chloride (ZD7288) (Sigma-Aldich), 10 μM 1-[2-[[(Diphenylmethylene)imino]oxy]ethyl]-1,2,5,6-tetrahydro-3-pyridinecarboxylic acid hydrochloride hydrochloride (NO711) (Sigma-Aldrich).

We performed Bayesian quantal analysis (BQA) by altering the extracellular Ca^2+^ and Mg^2+^ in two different conditions (Bhumbra & Beato, 2013a). One of the conditions were provide consistently a high release probability, in which the ACSF contained (in mM): 3 Ca^2+^ / 1.5 Mg^2+^. For the reduced release probability we tested two different composition (in mM): either 2 Ca^2+^ / 2 Mg^2+^ or 1.5 Ca^2+^/ 3 Mg^2+^. During BQA experiments the ACSF solution contained the following substances: 10 μM D-(−)-2-Amino-5-phosphonopentanoic acid (D-AP5) (Tocris), 10 μM 2,3-Dioxo-6-nitro-1,2,3,4-tetrahydrobenzo[*f*]quinoxaline-7-sulfonamide (NBQX) (Tocris). Each epoch of the BQA experiment contains a stable segment of 28 up to 42 unitary IPSCs (mean 32.75 ± 4.15). BQA experiments required at least 60 min of recording time (up to 90 minutes). We tested all epochs for possible long-term plasticity effect by measuring the linear correlation between IPSCs amplitude and elapsed time during the experiment, and we found no or negligible correlation (Pearson’s r values from all of the experiments (n= 8) were between −0.39 and 0.46, mean −0.01 ± 0.29).

### Immunohistochemistry and anatomical analysis

After electrophysiological recordings slices were fixed in a fixative containing 4% paraformaldehyde, 15% picric acid and 1.25% glutaraldehyde in 0.1 M phosphate buffer (PB; pH= 7.4) at 4°C for at least 12 hr. After several washes in 0.1M PB, slices were cryoprotected in 10% then 20% sucrose solution in 0.1 M PB. Slices were frozen in liquid nitrogen then thawed in PB, embedded in 10% gelatin and further sectioned into slices of 60 μm in thickness. Sections were incubated in a solution of conjugated avidin-biotin horseradish peroxidase (ABC; 1:100; Vector Labs) in Tris-buffered saline (TBS, pH= 7.4) at 4°C overnight. The enzyme reaction was revealed by 3’3-diaminobenzidine tetrahydrochloride (0.05%) as chromogen and 0.01% H_2_O_2_ as an oxidant. Sections were post-fixed with 1% OsO_4_ in 0.1 M PB. After several washes in distilled water, sections were stained in 1% uranyl acetate, dehydrated in ascending series of ethanol. Sections were infiltrated with epoxy resin (Durcupan (Sigma-Aldich)) overnight and embedded on glass slices. Three dimensional light microscopic reconstructions were carried out using Neurolucida system with a 100x objective.

### Surgery for imaging experiments

Experiments were conducted to the guidelines of University of Szeged Animal Care and Use Committee. Young adult (22 to 28 days old, (P) 24.75 ± 2.75) male Wistar rats were initially anaesthetized with halothane before urethane anaesthesia (1.4 g/kg of body weight) was administrated intraperitoneally. Body temperature was maintained at 37°C with a heating pad (Supertech Instruments, Pécs, Hungary). Before surgery dexamethasone sodium phosphate (2 mg/kg of body weight) was administrated subcutaneous, and carprofen (5 mg/kg of body weight) was administrated intraperitoneally. Anaesthetized animals head were stabilized in a stereotaxic frame and headbars were attached to the skull with dental cement (Sun Medical, Mariyama, Japan). Circular craniotomy (3 mm diameter) was made above the primary somatosensory cortex, centered at 1.5 mm posterior and 2.2 mm lateral from the bregma with a high-speed dental drill (Jinme Dental, Foshan, China). Dura mater was carefully removed surgically. Finally, the craniotomy was filled with 1.5 % agarose and covered with a coverslip to limit motion artifacts. The craniotomy was then submerged with HEPES buffered ACSF recording solution containing (in mM) 125 NaCl, 3.5 KCl, 10 HEPES, 1 MgSO_4_, 1 CaCl_2_, 0.5 d (+)-glucose, pH= 7.4.

### Two photon calcium imaging in L1

Before covering the craniotomy with the coverslip calcium indicator Oregon Green 488 BAPTA-1 AM (10mM) (OGB-1 AM, Thermo Fisher Scientific), and astrocytic marker sulforhodamine 101 (1 μM) (SR101, Thermo Fisher Scientific) were pressure-injected with a glass pipette (1-2 MΩ) in L1 cortical region under the visual guide of Zeiss Axio Examiner LSM7 (Carl Zeiss AG, Oberkochen, Germany) two-photon microscope using 40x water immersion objective (W-Plan, Carl Zeiss, Germany). Subsequently, the craniotomy was filled with agarose and covered with a coverslip. Imaging experiments were performed 1 hour after preparation. The activity of L1 interneurons was monitored during ipsilateral hindlimb electrical stimulation (Digimeter, Hertfordshire, United Kingdom, 200 mA, 10 ms). OGB-1 AM was excited at 800 nm wavelength with a femtosecond pulsing Ti:sapphire laser (Mai Tai DeepSee (Spectra-physics, Santa Clara, USA)). In the somatosensory hindlimb region, Z-stack image series (volume size 304 μm x 304 μm x 104 μm) were acquired. Calcium signals from interneurons were obtained within this volume in full-frame mode (256 x 100 pixel), acquired at a frequency of ~20 Hz. The Ca^2+^ dependent fluorescence change ΔF/F was calculated as R(t)=(F(t)-F_0_(t))/F_0_(t) based on Jia et al 2010 (Jia, Rochefort, Chen, & Konnerth, 2011). The R(t) denotes the relative change of fluorescence signal, F(t) denotes the mean fluorescence of a region of interest at a certain time point, F0(t) denotes the time-dependent baseline. Image stabilization was performed by ImageJ (Fiji) software using the Image stabilizer plugin (Kang Li, 2008; Schneider, Rasband, & Eliceiri, 2012). At the end of the experiments, few L1 neurons were filled with biocytin containing intracellular solution to make the immunohistochemical remapping easier.

### Tissue preparation for immunohistochemistry

After imaging experiments rats were deeply anaesthetized with ketamine and xylazine. Subsequently, perfusion was performed through the aorta, first with 0.9% saline for 1 min, then with an ice-cold fixative containing 4% paraformaldehyde in 0.1 M phosphate buffer (PB, pH= 7.4) for 15 min. The whole brain was extracted and stored in 4% paraformaldehyde for 24 hours, afterward in 0.1 M phosphate buffer (pH= 7.4) until slicing. Later 60 μm thick sections were cut from the same two-photon Ca^2+^ imaged brain area parallel to the pia mater and washed overnight in 0.1 M PB.

### Fluorescence immunohistochemistry and remapping

After several washes in 0.1 M PB, slices were cryoprotected with 10% then 20% sucrose solution in 0.1M PB than frozen in liquid nitrogen. The sections were incubated for two hours in Alexa-488 conjugated streptavidin (1:400, Molecular Probes) solved in Tris-buffered saline (TBS, 0.1 M; pH= 7.4) at room temperature to visualized the biocytin labeled cells. After several washes in TBS, sections were blocked in normal horse serum (NHS, 10%) made up in TBS, followed by incubation in mouse anti-Alfa-Actinin (1:20000, Sigma-Aldrich) diluted in TBS containing 2% NHS and 0.1% Triton X-100 at room temperature for 6 hours. Following several washes in TBS, Cy3 conjugated donkey anti-mouse (1:500, Jackson ImmunoResearch) secondary antibody was used to visualize the immunoreactions. After several washes in TBS then in 0.1 M PB, slices were counterstained with DAPI (4’, 6-diamidino-2-phenylindole, ThermoFisher Scientific). Sections were then mounted on slides in Vectashield (Vector Laboratories). Images were taken with LSM 880 confocal laser scanning microscope (Carl Zeiss AG, Oberkochen, Germany) using 40X oil-immersion objective (1.4 NA). Confocal image z-stack was tilted and panned manually to match with the *in vivo* two photon z-stack, allowing to profile imaged interneurons. During this process biocytin labeled neurons were used as a reference point.

### Data analysis

Electrophysiological data were analyzed with Fitmaster (HEKA Elektronik GmbH, Lambrecht, Germany), Origin 7.5 (OriginLab Corporation, Northampton, Massachusetts, USA), IgorPro (Wavemetrics, Portland, Oregon, USA). BQA experiments were analyzed using a Python written program (Bhumbra & Beato, 2013a), incorporating NumPy and SciPy packages. Two-photon calcium imaging data were acquired with ZEN 2 (Carl Zeiss AG, Oberkochen, Germany) and analyzed with MATLAB (The MathWorks, Natick, Massachusetts, USA), using Statistical Toolbox, Image Processing Toolbox, and custom written scripts.

### MCell model construction

The model framework was constructed in Blender v2.7. The simulation environment contained a 3D reconstruction of a dendritic structure based on a series section of electron microscopic data (available from VolRoverN program (Edwards et al., 2014)), and realistically positioned release sites of NGFCs. In the simulation environment the extracellular space was also modeled by creating an array of cubic cells containing cavities based on previous work from Tao et al (Tao et al., 2005). The cubic cells have 800 x 800 nm length, containing cavity that is 400 x 400 nm wide and 340 nm deep. The cubic cells and the dendritic segment were spaced 32 nm apart. The established array of cubic compartments creates an extracellular space that provides a volume fraction and tortuosity identical to the cortical brain tissue (volume fraction = 0.2 and tortuosity= 1.6). The overall dimensions of the modeled space surrounding the ultrastructurally reconstructed dendrite were 13.28 x 13.28 x 6.592 μm and the total volume was 1162.55 μm^3^.

Simulation of GABA_B_ receptor-GIRK channel interaction was carried out with MCell v3.4 (www.mcell.org)(Kerr et al., 2008). Custom Matlab scripts created the MDL (Model Description Language) file that required for MCell simulation. MCell simulated the release and diffusion of GABA, GABA_B_ receptors and GIRK channel interaction. First, to manage a biological like distribution for the receptors and channels a reaction cascade was used (Supplemetary Fig. 3). This cascade was constructed and tested in a simple simulation environment first, containing only a plane surface. At the beginning of every iteration, primary seed particles were placed on the dendritic membrane. Primary seed particles subdivide into secondary seed particles, that which then produce GABAB receptor or GIRK channel clusters. Those secondary seed particles that produce the GIRK channel clusters - which contain 1 to 4 channels - were immobile in the membrane. Meanwhile, the secondary seed particles that produce GABAB receptor clusters – which contain 1 to 8 receptors – can diffuse laterally in the membrane. At the end the distance was defined between the center of receptor and channel clusters by calculating the distance between each receptor and channel cluster in the two-dimensional plane surface. delay and the forward rate of the reaction was set to allow secondary seeds, that generate GABAB receptor clusters to diffuse to specified distance, resulting in the required GABA_B_ receptor-GIRK channel cluster distribution as seen from Kulik et al (A. Kulik, 2006). Optimization algorithm based on simulated annealing technique (Henderson, Jacobson, & Johnson, 2006; Kirkpatrick, Gelatt, & Vecchi, 1983) was written in Matlab for approximating the optimal values for the delay and the forward rate of the reaction. Optimal values of delay and the forward rate of the reaction was set to allow secondary seeds, that generate GABA_B_ receptor clusters to diffuse to specified distance, resulting in the required GABA_B_ receptor-GIRK channel cluster distribution as seen in Kulik et al. 2005.

Since we were interested in the interaction between the GABAB receptors and GIRK channels, our model does not include GABA_A_ receptors and GABA amino transporters. Previous work suggest that a single AP in the NGFC generates GABA concentration of 1 to 60 μM lasting for 20-200 ms (Karayannis et al., 2010), therefore in our model we used similar GABA concentration range of at 0.5 to 2 μm distance from the release sites.

Up to 6 MCell simulations were run with 1 μs time steps in parallel on pc with Intel (R) Core i7-4790 3.6 GHz CPU, 32 GB RAM. Total of 4278 iterations were simulated.

### Statistics

The number of experimental recordings used in each experiment is indicated in the text. Statistical tests were performed using Origin 7.5 (OriginLab Corporation, Northampton, Massachusetts, USA) and SPSS software (IBM, Armonk, NY, USA). Data are represented as mean ± standard deviation (SD). Data were first subject to a Shapiro-Wilk test of normality, and based on the result to the indicated parametric and non-parametric tests. Result were considered significantly different if p<0.05.

**Table 1.**
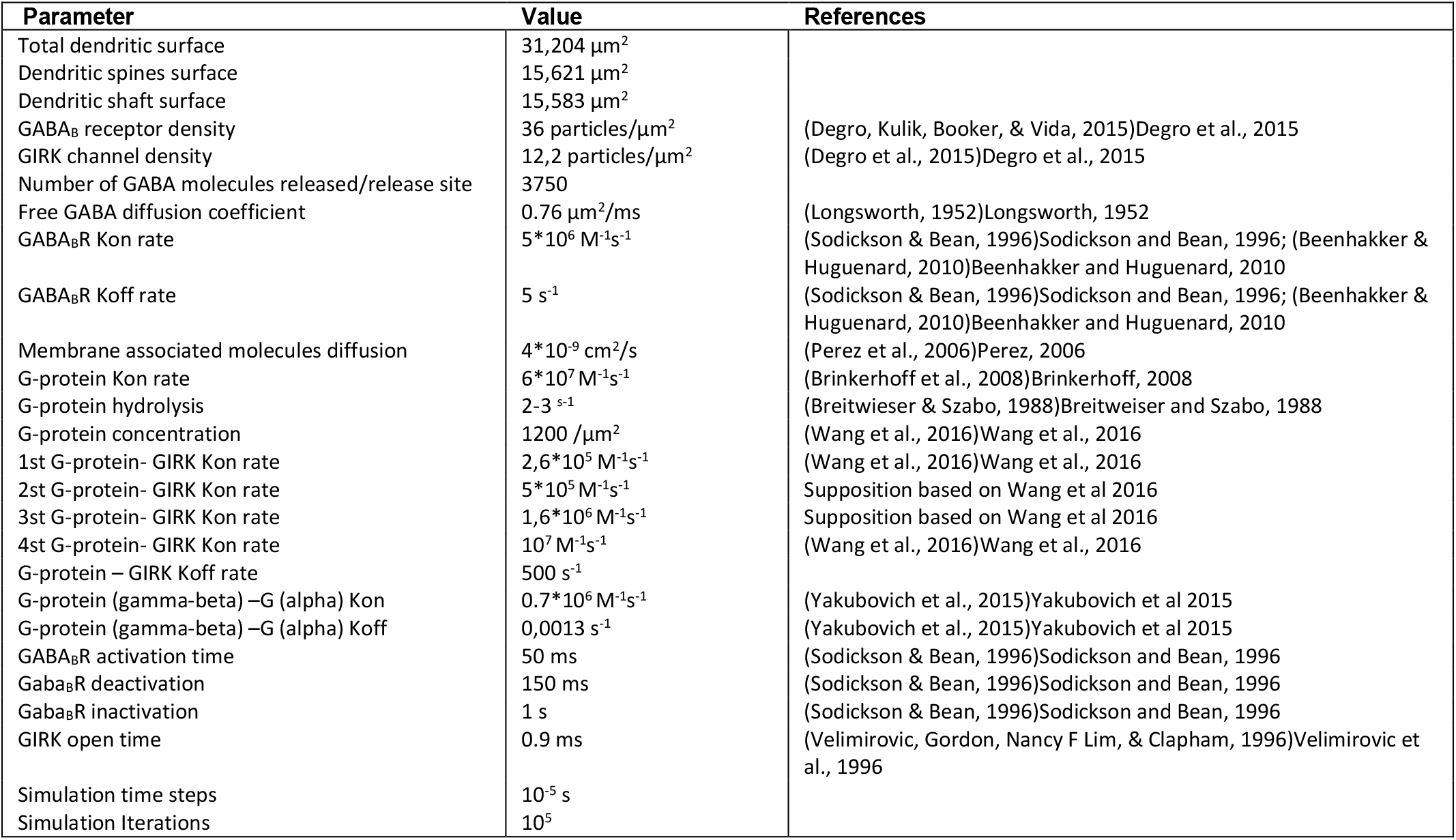
Parameters used for simulation

## Supplementary Information

**Sup. Fig 1.**
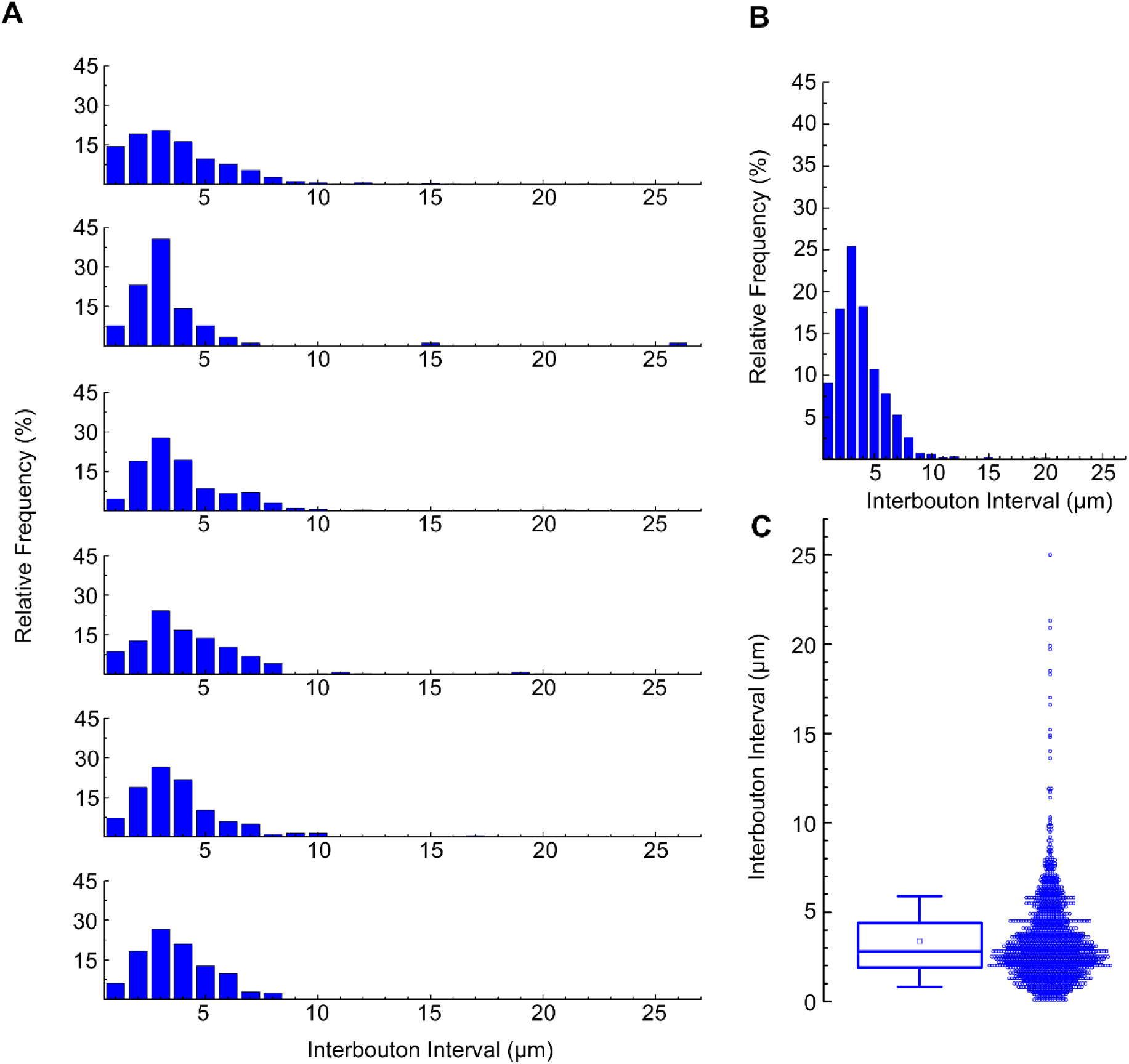
Related to Fig 2. Calculation of NGFC’s interbouton interval. (**a**) Distribution of interbouton distances from 6 NGFCs (number of boutons from each morphological analysis were n=442,91,213,252,291,207). The axonal segments were randomly selected (bin size = 1μm). (**b**) Average distribution of interbouton intervals in all NGFCs (bin size = 1μm). (**c**) Boxplot shows the mean±SD (3.357±2.535; n=1495) of all NGFCs’ interbouton interval.

**Sup. Fig 2.**
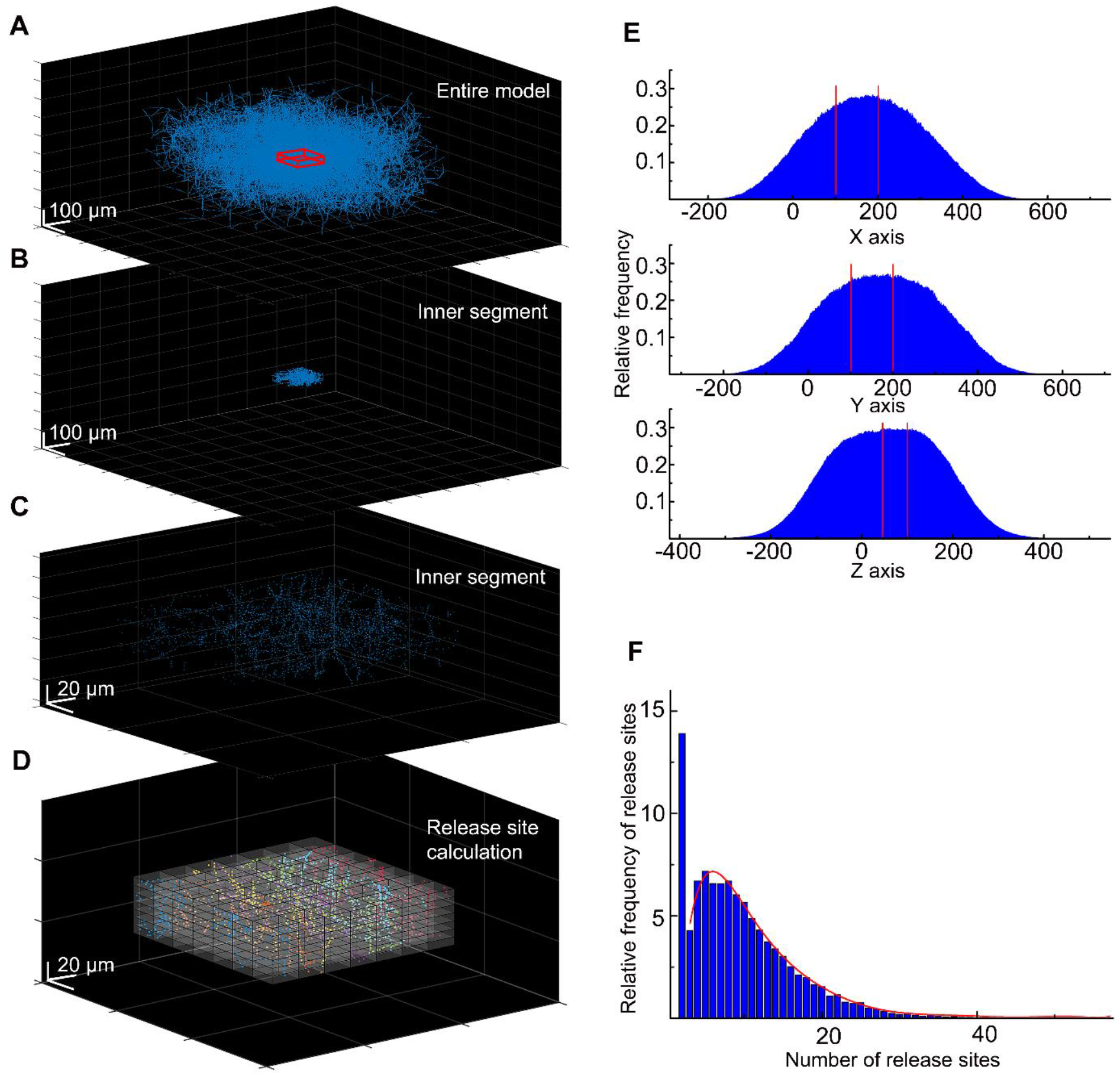
Related to Fig 6. Calculation of the NGFC release site density for the MCell model. (**a**) In the spatial model of the NGFC population, the density of axonal processes and release sites are rapidly declining around the model’s outskirts. To avoid underrepresentation of release sites, we collected data from the inner section of the model, from a 100×100×50 μm sized cuboid (showed in red). (**b**) Within these boundaries (the section between the two red lines) the high density of release sites is preserved alongside all three axes. (**c**) In the inner segment of the model, we calculated the frequency of release sites in smaller 13.28×13.28×3.592 μm sized cuboids. (**d**) We calculated the release site frequency within the inner segments in all (n=36) simulations in 13.28×13.28×3.592 μm volumes. (**e**) In each simulation, on average 2611.5±691.96 and a total of 94014 release sites were calculated. (**f**) Data shows that 13.9% of segmented volumes do not contain release sites. In 86.1% of segmented volumes, the release site frequency shows a skewed distribution from 1 to 56 (mean:7.61±6.71). The red line shows the fitting of this data.

**Sup. Fig 3.**
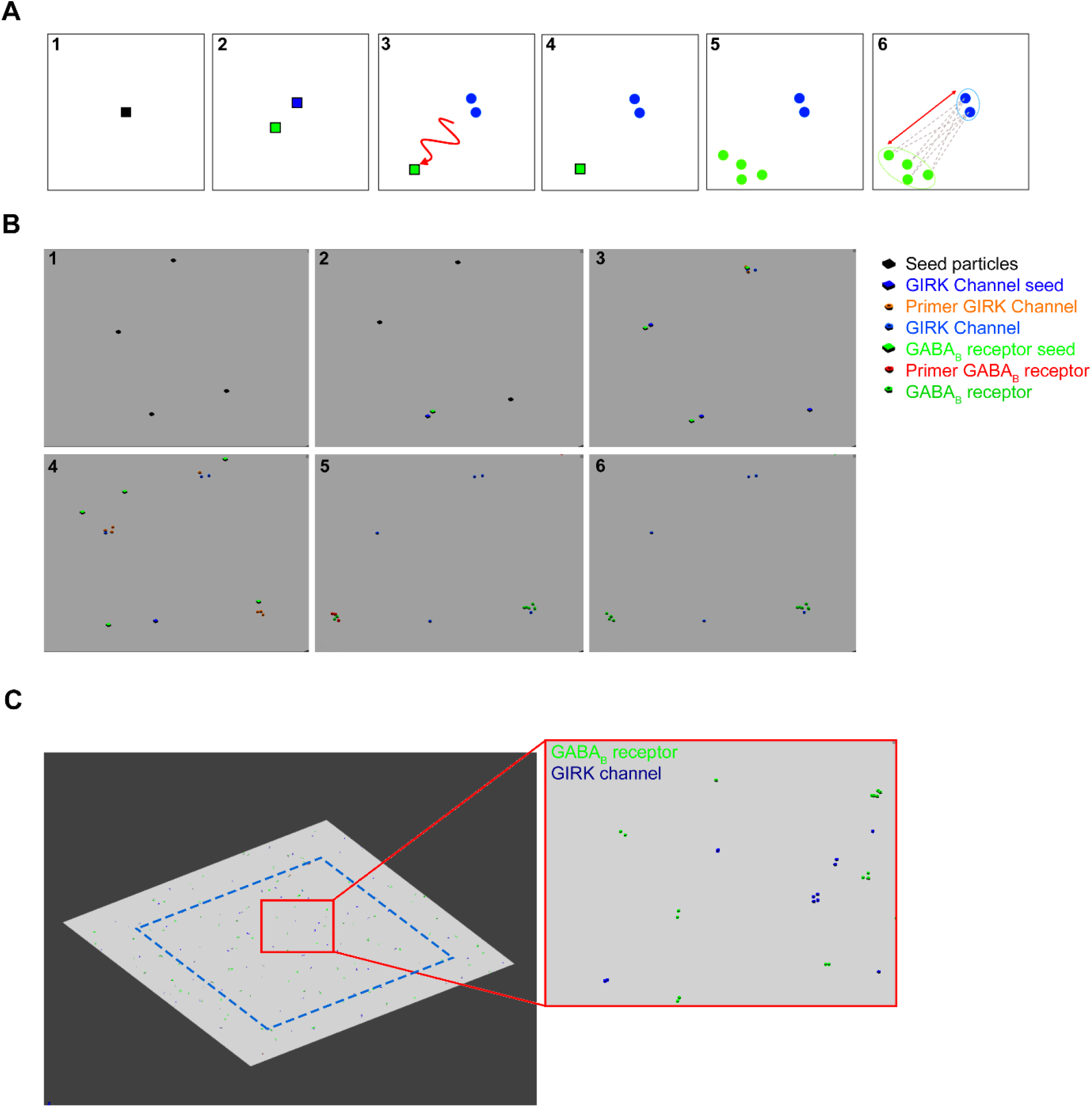
GABA_B_ receptor and GIRK channel distribution, created by a cascade reaction. (**a**) To create a pseudorandom distribution the molecular interactions the following steps were defined as the schematic concept: (1) Seed particles (black square) were placed randomly along the membrane surface. (2) The seed particles produce two distinctive particles: the first particle, the GIRK channel seed (blue square) that immediately creates GIRK channels (blue circles) and it remains immobile. (4) The second particle is the GABA_B_ receptor seed (green square), which has the ability to laterally diffuse. (5) The probabilistic distance between the static GIRK channels and the GABA_B_ receptor seed can be adjusted by limiting the time while it is mobile. After became immobile, the GABA_B_ receptor seed produces GABA_B_ receptors. (6) The distance between each GIRK channel and the nearest GABA_B_ receptors was calculated. (**b**) Image sequences show the particle interactions in the MCell model: (1) Primer seed particles were placed randomly, (2) which produce GIRK channel seed and GABA_B_ receptor seed particles. (3) GIRK channel seed particles remain in the same place, the GABA_B_ receptor seed particles laterally diffuse away. (4) GIRK channel seed particles produce 1 to 4 GIRK channels by first creating 4 primer GIRK channel particles that either disappear or produce a final GIRK channel. This step was created to avoid uniform cluster generation. (5) GABA_B_ receptor seeds lateral diffusion time was optimized to regulate the probabilistic distance between the cluster. GABA_B_ receptor seed generated 1 to 8 GABA_B_ receptors, that either disappeared or produced a functional receptor. (6) Finally, distances were measured between the closest receptor and channel cluster. (**c**) Images show the two-dimensional plane and inset of a zoomed image of the final distribution. Distances were only measured from the GIRK channels that were inside the blue dashed line.

**Sup. Fig 4.**
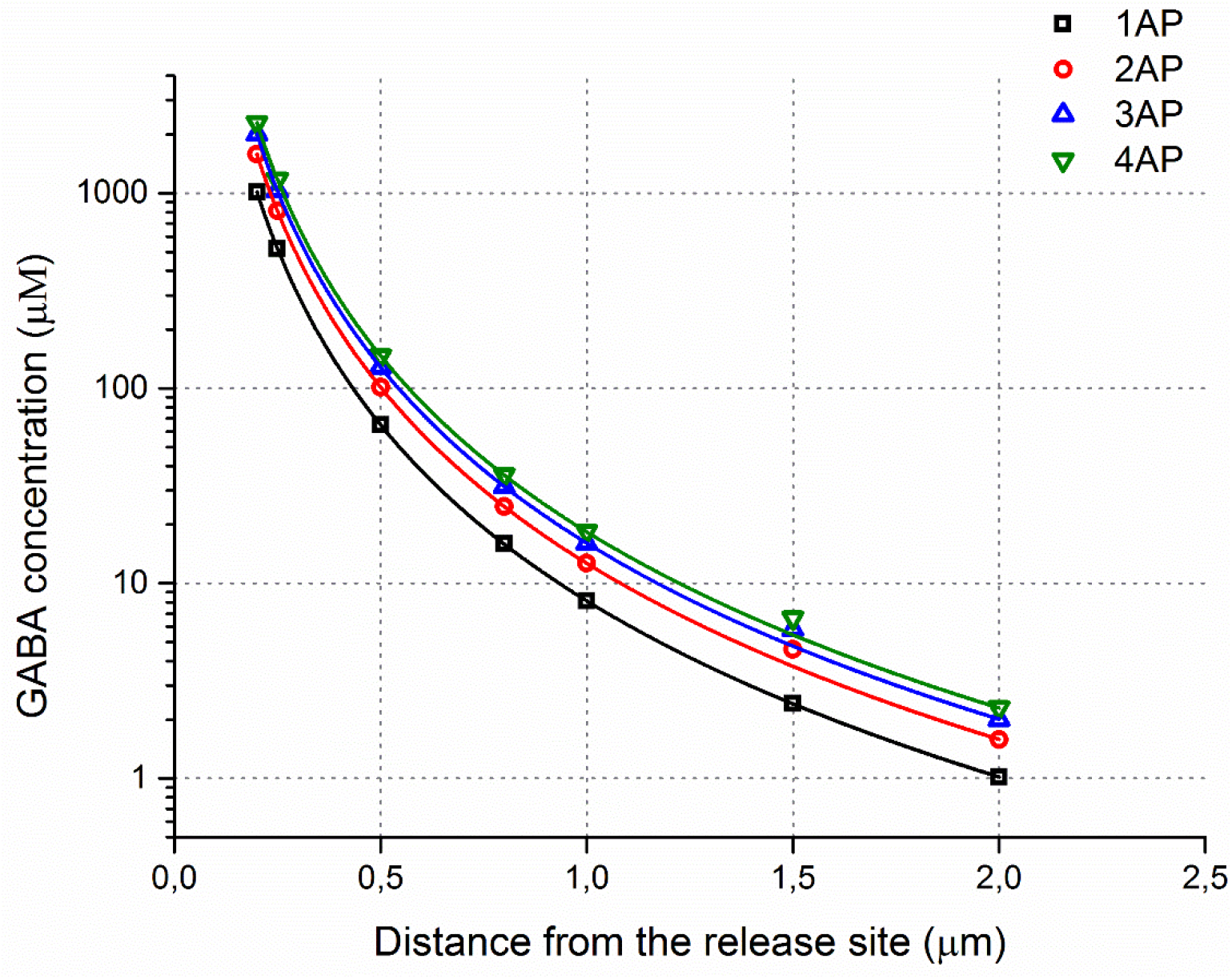
Estimated GABA spatial concentration profiles during multiple releases. A plot of the GABA concentrations versus distance from the release site in the MCell model.

**Sup. Table 1.**
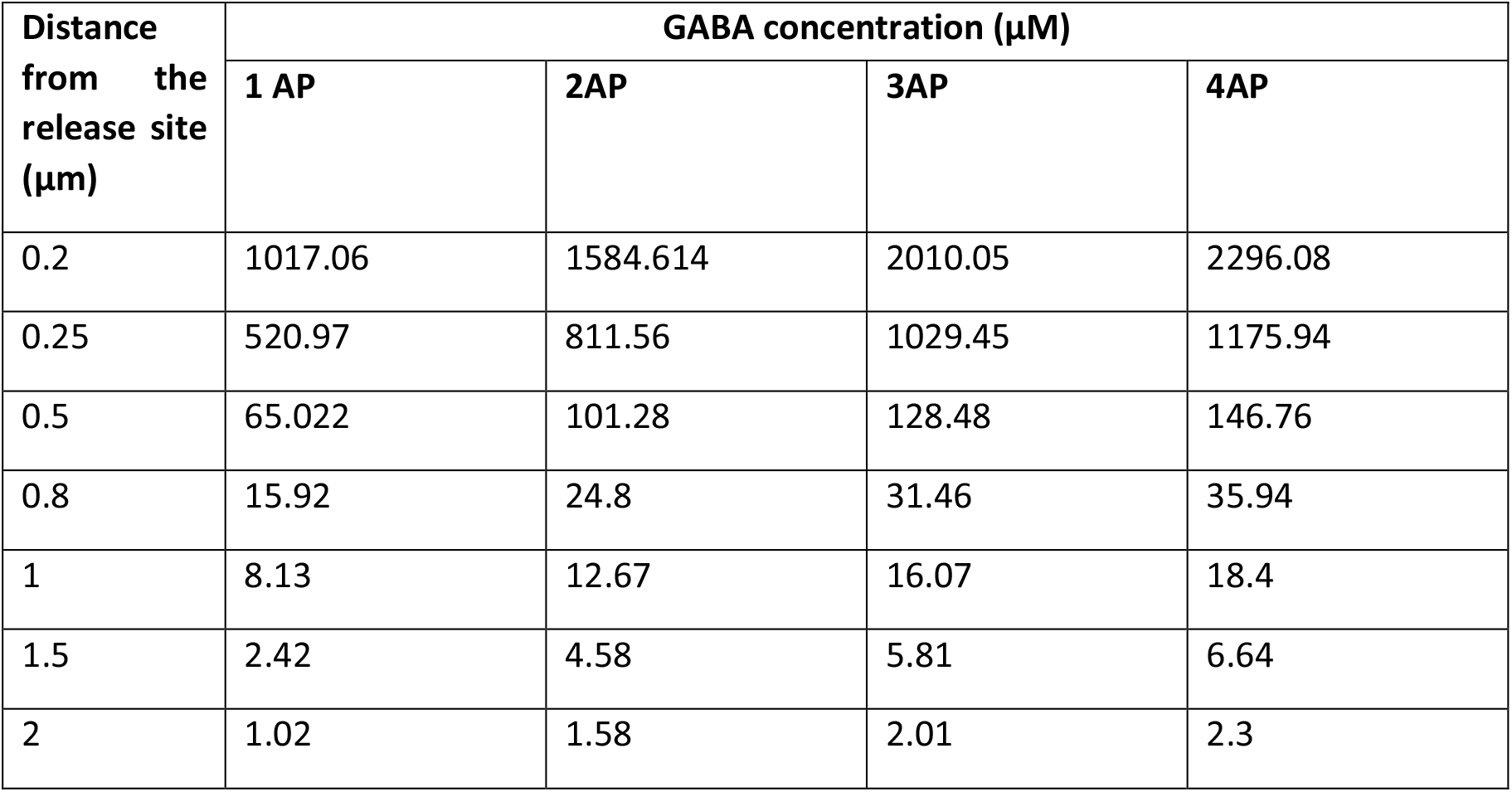
Different estimated GABA concentrations as a function of distance.

